# The covariance environment defines cellular niches for spatial inference

**DOI:** 10.1101/2023.04.18.537375

**Authors:** Doron Haviv, Mohamed Gatie, Anna-Katerina Hadjantonakis, Tal Nawy, Dana Pe’er

## Abstract

The tsunami of new multiplexed spatial profiling technologies has opened a range of computational challenges focused on leveraging these powerful data for biological discovery. A key challenge underlying computation is a suitable representation for features of cellular niches. Here, we develop the covariance environment (COVET), a representation that can capture the rich, continuous multivariate nature of cellular niches by capturing the gene-gene covariate structure across cells in the niche, which can reflect the cell-cell communication between them. We define a principled optimal transport-based distance metric between COVET niches and develop a computationally efficient approximation to this metric that can scale to millions of cells. Using COVET to encode spatial context, we develop environmental variational inference (ENVI), a conditional variational autoencoder that jointly embeds spatial and single-cell RNA-seq data into a latent space. Two distinct decoders either impute gene expression across spatial modality, or project spatial information onto dissociated single-cell data. We show that ENVI is not only superior in the imputation of gene expression but is also able to infer spatial context to disassociated single-cell genomics data.

## INTRODUCTION

Intense interest in cellular interactions and tissue context has spurred the growth of multiplexed spatial transcriptomics and antibody-based technologies and datasets, sparking the need for computational approaches to identify biological patterns within tissues^1–5^. The local neighborhood, or niche, of a cell is a useful resolution for defining cell interactions; it may represent functional anatomical subunits (such as stem cell niches) and is a basis for identifying larger spatial patterns. However, we lack efficient representations of the cellular microenvironment that retain the full richness of the data and can be used to effectively compare niches and their distributions. At the same time, we need to address the limited molecular plexity of high-resolution spatial profiling technologies.

Most methods for analyzing spatial data characterize each niche by tabulating discrete cell types within a given region or radius^6–9^. While these have generated important discoveries, including differences between healthy and diseased tissue^2, 6^, they were developed for low-plex antibody-based imaging methods (∼30 markers) that devote most markers to cell typing. Spatial transcriptomics methods, including commercial platforms, can now profile hundreds of genes^10–16^, meaning that analysis at the cell type level leads to substantial information loss. In single-cell genomics, the switch from discrete cell-typing to continuous approaches such as diffusion maps^17^ and pseudotime^18, 19^ has driven remarkable discovery. Moreover, setting thresholds for continuous cellular phenotypes is subjective and invokes problems of instability and bias. Even within highly discrete cell types, vast and meaningful variation often exists, such as the spectrum of activated and metabolic states within immune cell types^20–22^.

We thus need a niche representation that considers the full expression matrix and its continuous nature, and that enables robust and efficient comparisons. However, most approaches are based on cell typing, which compresses gene expression to one-hot encodings of cell type labels, thus losing information about cell-cell similarity and complex patterns of gene expression, including covariation in genes that are coregulated across cell states. Here, we develop the covariance environment (COVET), a compact representation of a cell’s niche which assumes that interactions between the cell and its environment create biologically meaningful covariate structure in gene expression between cells of the niche. We develop a corresponding distance metric that unlocks the ability to compare and analyze niches using the full toolkit of approaches currently employed for cellular phenotypes, including dimensionality reduction and clustering.

Imaging-based spatial transcriptomics technologies face issues such as signal crowding, tissue degradation and drift, cumulative decoding error and noise, and long collection times, which practically limit quantification to hundreds of genes. Methods have been developed that impute spatial information for genes not measured in the spatial modality, by integrating matched, transcriptome-wide single-cell RNA sequencing (scRNA-seq) data^7, 23, 24^. Yet almost no integration methods explicitly model cellular microenvironment context from the spatial data, arguably due to not having a reliable representation of spatial context like COVET. Instead, they only carry out partial integration, by modeling the expression of just those genes measured in the spatial modality, thereby limiting inference power.

To achieve transcriptome-wide spatial inference, we developed environmental variational inference (ENVI), a conditional variational autoencoder (CVAE)^25, 26^ that simultaneously incorporates scRNA-seq and spatial data into a single embedding. ENVI leverages the covariate structure of COVET as a representation of cell microenvironment and achieves total integration by encoding both genome-wide expression and spatial context (the ability to reconstruct COVET matrices) into its latent embedding. We demonstrate that our approach is effective on data from a variety of multiplexed spatial technologies, and that it outperforms other methods in accurately imputing the expression of genes in diverse developmental contexts. We also show that ENVI can be used to project valuable spatial information onto dissociated scRNA-seq data, and that it can capture and express continuous variation along spatial axes across large complex tissue regions.

## RESULTS

### The covariance environment defines spatial neighborhoods

To move beyond cell-type fraction and characterize niches in a manner that leverages all measured genes and enables quantitative comparison, we developed the COVET framework. Our core assumption is that a cell affects—and is also affected by—cells in its vicinity, generating covarying patterns of expression among the interacting cells. Our framework includes three components: 1) COVET, a robust per-cell representation of neighborhood information based on modified gene-gene covariance among niche cells, 2) a distance metric that is essential for comparing and interpreting niches, and 3) an algorithm to efficiently compute distance. Unlike mean expression, gene-gene covariance captures the relationships among genes and cell states that are shaped by cellular interplay within the niche. These relationships are rich, stable, and enriched for biological signal; moreover, they contain substantial hidden information from unmeasured genes, providing an advantage for imputation tasks.

To calculate COVET, we first define the niche of each cell in a dataset by the *k* spatial nearest neighbors of that cell, and then compute each niche’s gene-gene shifted covariance matrix (**Fig. 1a** and Methods). Shifted covariance is a modification of the classic covariance formulation, in which we use mean expression across the entire dataset rather than local mean expression as a reference. This constructs each cell’s covariance matrix relative to the entire population, and critically enables direct comparison between niches, highlighting their shared and unique features. Gene-gene covariance provides the additional benefit of being more robust to technical artifacts^21^, facilitating data integration and niche comparison across technologies.

**Figure 1.**
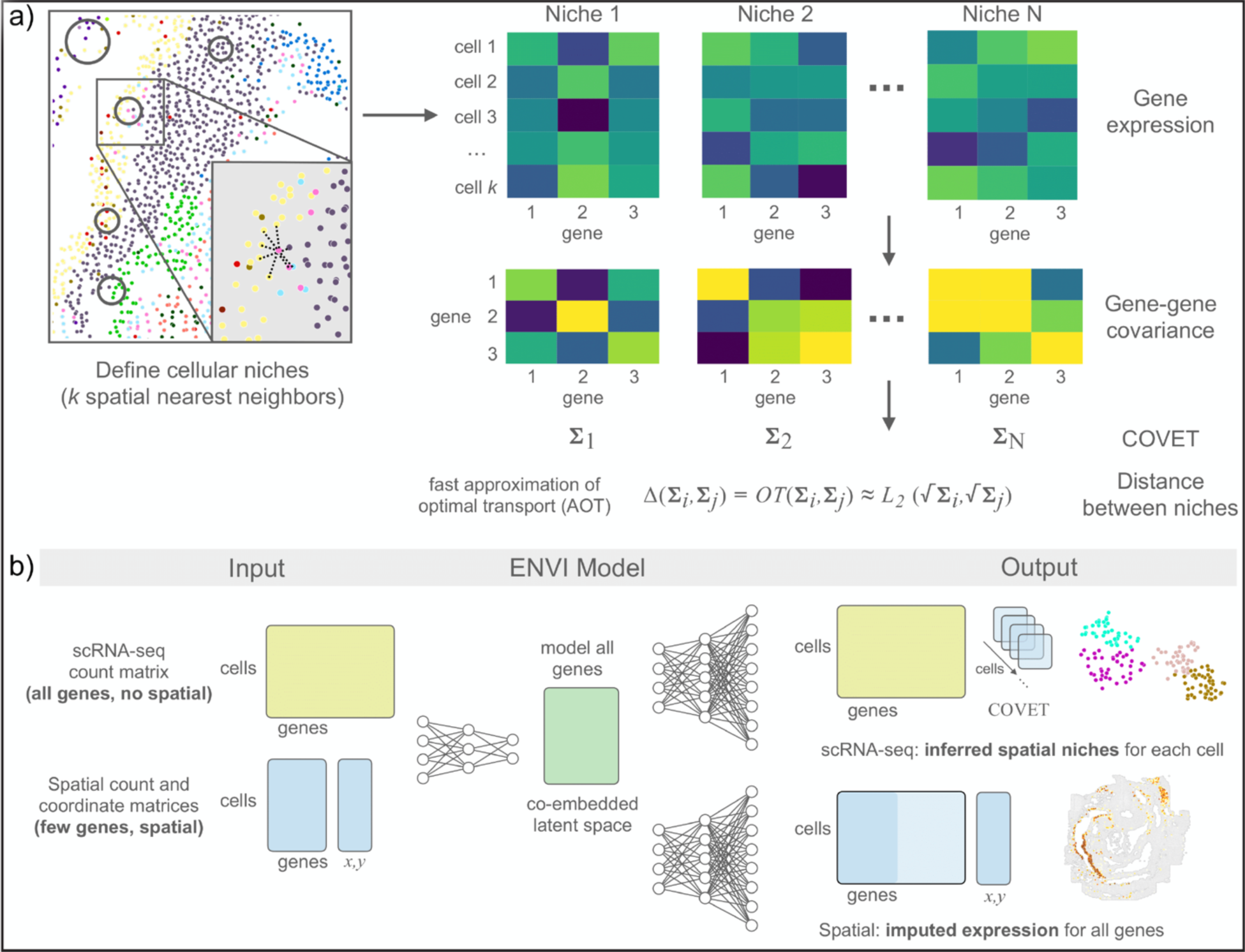
A covariance-based framework for characterizing spatial niches powers ENVI integration of single-cell and spatial data for robust transcriptome-wide spatial inference. **a**, Schematics indicate steps in spatial covariance computation and ENVI operation. Each covariance environment (COVET) matrix characterizes a cell’s niche, comprised of itself and *k* nearest spatial neighbors, based on the shifted covariance of gene expression within the niche. Shifted covariance is calculated relative to mean expression in the sample, enabling meaningful comparison of niches. Distance between niches is determined by an efficient approximation of optimal transport. **b**, ENVI is a conditional autoencoder that simultaneously embeds scRNA-seq and multiplexed spatial transcriptomic data into a unified latent embedding. ENVI models all genes (including those not imaged with spatial transcriptomics) and uses the COVET framework to represent information about cellular environment. Two decoder networks allow ENVI to project spatial information onto single-cell data, and to impute spatial expression transcriptome-wide.

Despite being a compact and powerful representation of the niche, COVET requires a metric for comparison to be interpretable. Niche similarity cannot be determined by simply subtracting the cell-by-gene expression of matrices of two niches, since the result depends on cell order, which is set arbitrarily (it will change if an image is rotated, for example). We thus seek to quantify niche similarity in a permutation-invariant manner, for which the Fréchet distance provides a closed-form solution^27^. However, calculating Fréchet distance is computationally demanding, so we developed an approximation (approximate optimal transport, AOT) that reduces runtime by over an order of magnitude, and is substantially faster than another common metric, the Bhattacharyya distance^28^ (**Supplementary Fig. 1a**). AOT yields similar results to true optimal transport and Bhattacharyya distance, and a GPU implementation takes under 1 minute to compute the cell-cell AOT distance matrix of 100,000 cells (**Supplementary Fig. 1b,c**).

As AOT can be computed via Euclidean distance, which underlies many standard single-cell analyses such as clustering^29^, diffusion components^30^ and uniform manifold approximation and projection (UMAP)^31^, niches can now be analyzed with the same algorithms designed to analyze phenotypes. Clustering niches can characterize canonical environments, visualization can be used to observe their relationships, and trajectory analysis can capture continuous trends, enabling facile interpretation. COVET thus provides a robust, tractable, and computationally efficient representation of cellular niches, which can uncover continuous trends of interplay between a cell and its microenvironment, derived from a mathematically principled formulation based on optimal transport.

### The ENVI algorithm

ENVI employs a conditional variational autoencoder to infer spatial context in scRNA-seq data and to impute missing genes in spatial data, by mapping both modalities to a common embedding (**Fig. 1b** and Methods). Unlike other CVAEs used for spatial inference^23, 32, 33^, which only model genes measured in both modalities, ENVI explicitly models spatial information and gene expression genome-wide. More importantly, it uses the COVET matrix to represent spatial information, and simultaneously trains on samples from both spatial and single-cell datasets, optimizing a single latent space to decode the full transcriptome and spatial context for both modalities.

ENVI architecture includes a single encoder for both spatial and single-cell genomics data, and two decoder networks—one for the full transcriptome, and the second for the COVET matrix, providing spatial context. The requisite for decoding the spatial niche (and the use of a second decoder) is a unique aspect of ENVI. Intuitively, ENVI uses gene expression in the cell paired with its niche information (COVET) to learn an ‘environment’ regression model, which infers spatial context from gene expression input, and simultaneously, an ‘imputation’ regression model trained to reproduce the full scRNA-seq dataset from the gene subset profiled by spatial transcriptomics. The nonlinear network architecture can capture complex dependencies between the variables.

Sequencing and spatial technologies measure different parameters, in addition to harboring technical differences that result in different data distributions and dynamic ranges (**Supplementary Fig. 2a**). We designed ENVI to take these into account by marginalizing technology-specific effects on expression, augmenting the standard VAE by adding an auxiliary binary neuron to the input layers of encoding and decoding networks for each modality. Moreover, ENVI parameterizes each modality with different probabilistic distributions, modeling single-cell data with a negative binomial by default to account for drop-out^34^, and spatial data with a Poisson by default to reflect the high capture rate of FISH-based technologies^3^. ENVI thus integrates, imputes and reconstructs spatial context with a single end-to-end model, utilizing deep learning for high-dimensional regression and rare signal extraction, and variational inference for optimal integration of scRNA-seq and spatial data. The method scales to atlas-size datasets including millions of cells with constant time computational complexity (**Supplementary Fig. 2b**).

### ENVI imputes complex spatial patterns underlying mouse gastrulation

To demonstrate ENVI’s abilities, we analyzed a 350-gene sequential fluorescence in situ hybridization (seqFISH)^35^ and scRNA-seq dataset^36^ of mouse organogenesis at embryonic day 8.75 (E8.75) (**Fig. 2a**). The developing embryo undergoes rapid cell proliferation, differentiation and movement to create stereotypical but highly complex multi-lineage cellular patterns and spatial gradients, unlike the discrete organized layers of adult brain tissue^3, 37, 38^ that dominate current spatial transcriptomic datasets. It thus presents a particularly challenging context to evaluate data integration and gene imputation. To successfully transfer information between technologies, spatial and scRNA-seq datasets must co-embed in the latent space, as neighboring cells in the latent space share similar decodings. The most basic evaluation of any embedding-based data integration method, therefore, is whether data across the two technologies co-embed successfully. Despite the complexity of seqFISH and scRNA-seq data and the technical differences between them, the embedding learned by ENVI trained with default parameters correctly maps major cell types to the combined latent space (**Fig. 2b**).

**Figure 2.**
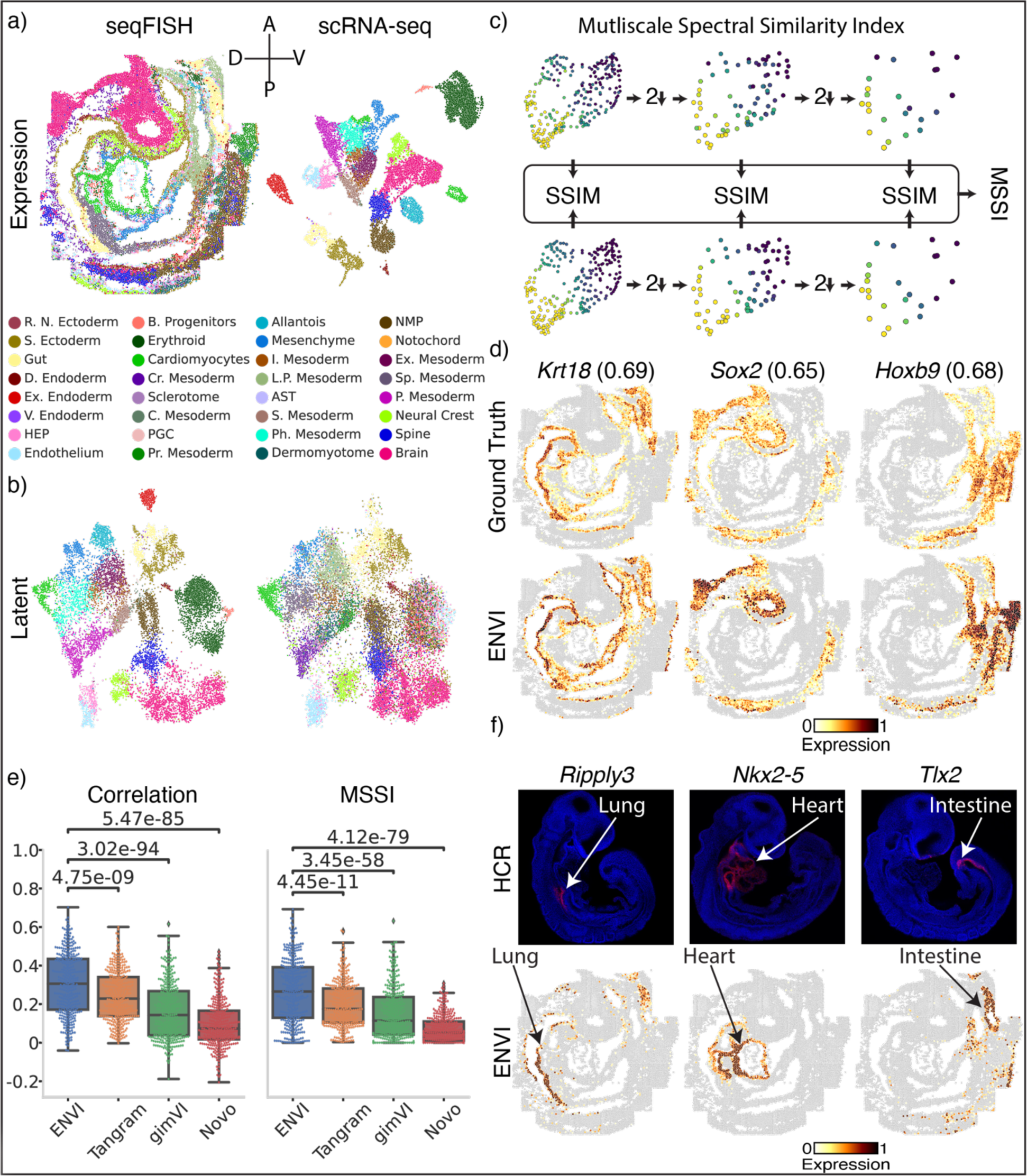
ENVI accurately recovers the expression of embryonic genes not imaged by multiplexed mRNA in situ hybridization. **a**, seqFISH^35^ image of an E8.75 mouse embryo sagittal section (left) and uniform manifold approximation and projection (UMAP) embedding of matched scRNA-seq data^36^ (right), both colored by major cell type compartment. **b**, UMAPs of ENVI latent embedding learned from mouse embryo data. Cells from seqFISH (left) and scRNA-seq (right) data are colored as in **a**. **c**, Schematic of multiscale spectral similarity index (MSSI) computation (Methods), for comparing two spatial expression profiles. Each profile is iteratively down sampled using spectral pooling on the cell proximity graph, and the standard structural similarity index metric (SSIM) is computed at each scale. MSSI is a weighted geometric mean of the SSIM computed at 5 scales, providing a spatially aware similarity metric on a scale from 0 to 1. **d**, seqFISH measurement in log counts (ground truth) and ENVI imputation for 3 withheld genes, marking endoderm (*Krt18*), neural stem (*Sox2*) and mesoderm (*Hand1*). MSSI values appear in parentheses for each gene. **e**, Pearson correlation and MSSI scores between log of seqFISH and imputed expression across all genes predicted from fivefold cross-validation, comparing four algorithms run with default parameters (Methods). Novo, NovoSpaRc. Boxes and lines represent interquartile range (IQR) and median, respectively; whiskers represent ±1.5 x IQR; values between box pairs indicate significance, one-sided relative t-test. **f**, ENVI imputation (bottom) of organ marker genes not profiled by seqFISH: *Ripply3* (lung), *Nkx2-5* (heart) and *Tlx2* (intestine). Imputed expression matches associated organs and is validated by whole-mount HCR in situ (top). R. N. Ectoderm, Rostral neurectoderm; S. Ectoderm, Surface Ectoderm; D., Definitive; V., Visceral; Cr., Cranial; C, Caudal; Pr., Presomitic; I., Intermediate; L.P., Lateral Plate; S. Mesoderm, Somitic Mesoderm; Ph., Pharyngeal; Ex., Extraembryonic; Sp., Sphlanic; P., Paraxial; HEP, Haematoendothelial progenitors; B., Blood; PGC, Primordial germ cells; AST, Anterior somitic tissue; NMP, Neuromesodermal Progenitors.

Current fluorescent in situ hybridization (FISH)-based technologies only image the patterns of expression of dozens to hundreds of genes^10, 35, 38^, prompting the development of algorithms to impute the spatial patterns of unmeasured genes^7, 23, 24, 39, 40^. We sought to evaluate ENVI’s ability to impute spatial gene expression. Previous work^7, 23, 41^ used Pearson correlation and mean squared error between imputed and ground-truth expression to evaluate the quality of imputation. However, both metrics are computed on a per-cell basis, without any awareness of cell-cell proximity or other spatial features. To evaluate concordance between spatial patterns, we developed the multiscale spectral similarity index (MSSI), a metric that can capture similarity between spatial patterns by taking cell-cell proximity into account (**Fig. 2c** and Methods). MSSI borrows from the multiscale structure similarity index measure (MS-SSIM)^42^, a spatial pattern similarity metric widely used in computer vision that iteratively subsamples the image and assesses similarity at multiple resolutions. Our MSSI metric uses a cell-cell neighbor graph based on spatial proximity to generate a series of images at progressively lower resolutions by aggregating spatially proximal cells, then applying SSIM to compare similarity at each resolution. MSSI thus provides a spatially aware similarity metric that uses full count matrices and incorporates patterning at the cellular level rather than pixel level. While we use MSSI to benchmark imputation, it can be applied more generally to compare similarity between the expression of different genes^43^, and other use cases.

We implemented a five-fold cross-validation scheme to generate ground truth (Methods) and compared ENVI imputation of held-out genes to this ground truth using both MSSI and Pearson correlation. As representative examples, we chose genes with clear spatial expression in endoderm (*Krt18*), neural stem (*Sox2*) and mesoderm (*Hand1*). The imputed expression of each gene was not only visually similar to ground truth (**Fig. 2d**), but also matched biological expectation in the correct organ. We also found that genes with correctly predicted organ-specific expression often have high MSSI scores but low Pearson correlations, supporting the importance of a spatially aware metric (**Supplementary Fig. 3a**).

We compared ENVI against widely used methods, including Tangram^7^ and gimVI^23^, which were recently shown to outperform other integration methods^41^. We also included NovoSpaRc^24^, because it uses fused optimal transport to explicitly model spatial context as the in-situ distance between samples. ENVI significantly outperforms all other methods based on both MSSI and Pearson correlation (**Fig. 2e**).

Finally, we evaluated ENVI’s ability to impute genes beyond those in the 350-gene panel by assessing canonical markers of the developing lung (*Ripply3*)^44^, heart (*Nkx2-5*)^45^ and intestine (*Tlx2*)^46^. In situ hybridization chain reaction (HCR) imaging^47^ of each marker further validated organ-specific expression at E8.75, before these organs form. Indeed, ENVI imputation and HCR imaging both localized each gene specifically to their respective organs: *Ripply3* to dorsal and ventral lung, *Nkx2-5* to the heart in the central embryo, and *Tlx2* to the posterior gut region (**Fig. 2f**). By contrast, Tangram and gimVI predicted weaker expression in the relevant organ and anomalous expression beyond the organ (**Supplementary Fig. 3b**).

### ENVI accurately imputes expression across datasets and modalities

To systematically evaluate the quality of ENVI imputation across a range of technologies and tissues, we assessed 33-gene osmFISH^3^, 42-gene ExSeq^37^ and 252-gene MERFISH^38^ datasets from murine somatosensory cortex, visual cortex, and primary motor cortex, respectively, using withheld genes to benchmark spatial imputation as for the gastrulation dataset (**Fig. 2e**).

We chose the osmFISH dataset, which was generated by iterative single molecule FISH without barcoding or amplification^3^, to assess whether transcriptome-scale inference is possible from only 33 imaged genes. The osmFISH data consists of a single image containing 4,530 cells, and 3,005 matched scRNA-seq cells after processing (**Supplementary Fig. 4a**). Due to the limited sample size, combining both technologies into a unified embedding is challenging^41^. Thus, we first evaluated whether the same cell types align and different cell types are separated in our latent embedding (**Supplementary Fig. 4b**). One other method in our comparison, gimVI, is also an autoencoder and learns a latent embedding during integration. We found that ENVI outperforms gimVI both by visual inspection (**Supplementary Fig. 4b,c**) and by explicitly quantifying distances between cell type latent embeddings across technologies (**Supplementary Fig. 4d** and Methods).

Using a leave-one-out approach to evaluate spatial gene imputation, we tested all 33 genes and found that ENVI substantially outperforms alternative methods (**Fig. 3a**). To determine whether ENVI can impute genes that were not imaged, we leveraged Allen Brain Atlas ground-truth data for mouse brain cortex (mouse.brain-map.org) and confirmed that ENVI correctly imputes layer-specific spatial expression for *Dti4l*, *Rprm* and *Ntst4* in the L2/3, L5/6 and CA1 regions, respectively (**Fig. 3b**).

**Figure 3:**
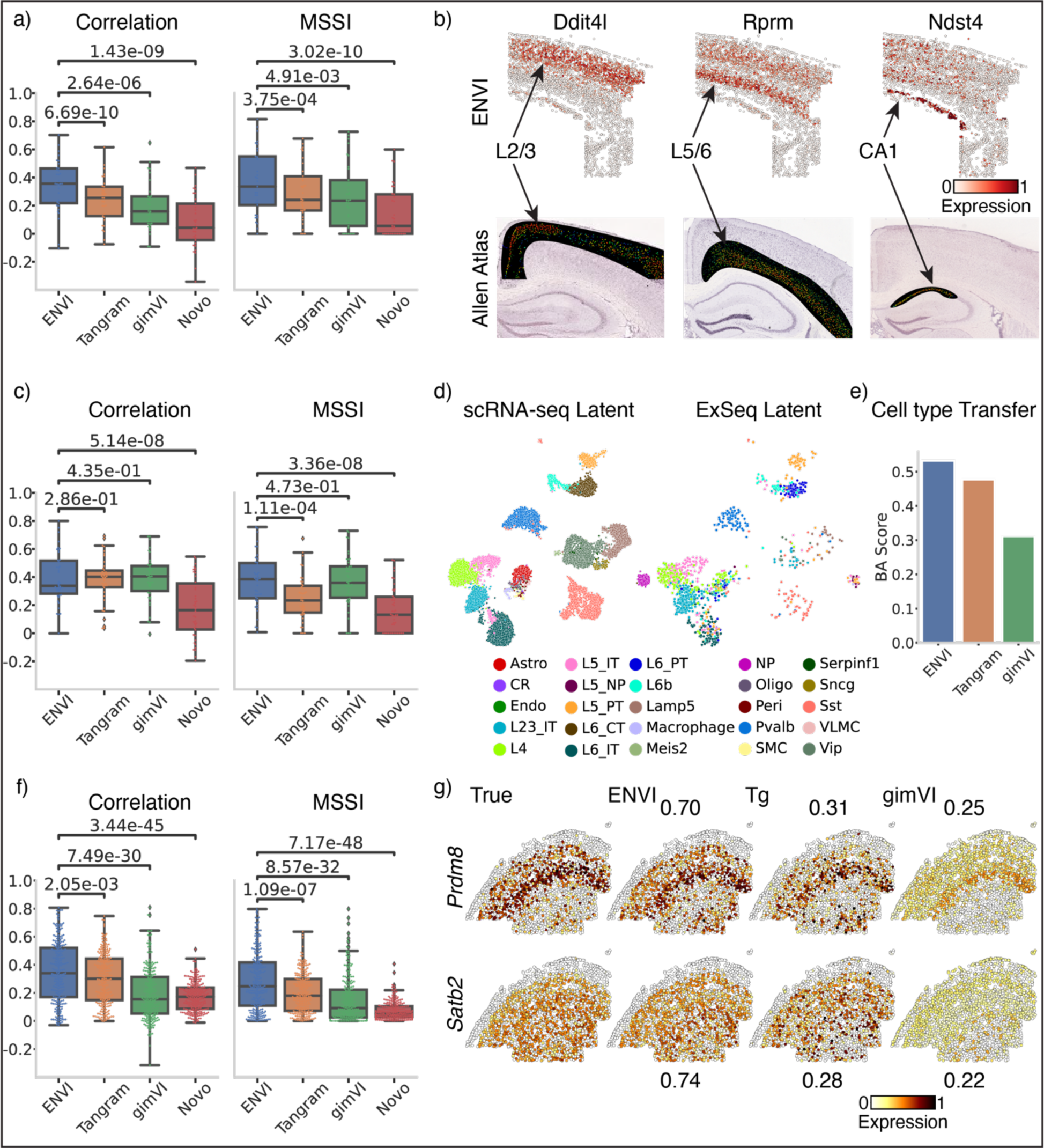
ENVI outperforms alternative methods on data from different imaging technologies and tissue contexts. **a**, Benchmarking of imputation based on leave-one-out training, evaluated by Pearson correlation and MSSI on a 33-gene osmFISH dataset of the somatosensory cortex^3^. **b**, ENVI-imputed expression of unimaged cortical markers *Ddit4l* (L2/3), *Rprm* (L5/6) and *Ndst* (Hippocampus, CA1) (top) and corresponding expression in the Allen Brain Atlas (mouse.brain-map.org) (bottom). **c**, Leave-one-out benchmarking of imputation based on a 41-gene ExSeq imaging dataset of the visual cortex^37^. **d**, ENVI latent embeddings of ExSeq and scRNA-seq data from the visual cortex. UMAP visualization colored by cell type, labeled according to Alon et al.^37^. **e**, Quality of data integration, based on balanced accuracy (BA) score between true and predicted classification from transferring cell-type labels from scRNA-seq to ExSeq data. **f**, Five-fold cross-validation of imputation based on a 252-gene MERFISH dataset from the primary motor cortex^38^. **g**, Expression of *Prdm8* (top) and *Satb2* (bottom) by MERFISH or reconstructions. Tg, Tangram. Numerical values represent MSSI scores compared to ground truth. **a**,**c**,**f**, Boxes and lines represent IQR and median, respectively; whiskers represent ±1.5 x IQR; *P*-values above box pairs, one-sided relative t-test. Novo, NovoSpaRc.

We next assessed how well ENVI extends to imaging technologies that are not based on FISH. ExSeq uses in situ sequencing to image individual RNA genes and is the basis of the 10x Genomics Xenium imaging platform. We validated ENVI on a 42-gene ExSeq assay from the visual cortex (1,271 cells) and a matched scRNA-seq dataset (13,586 cells)^37^, which together identify 25 populations comprising both neuronal and non-neuronal cell types (**Supplementary Fig. 5a**). Beyond representing a new technology, ExSeq data is formidable for its sparsity (possibly due to the use of expansion microscopy), which significantly hampers the performance of spatial and scRNA-seq integration methods^41^; some cell types in the visual cortex, especially from the inhibitory neuron compartment, were scarcely present (for example, only 13 *Vip* and 25 *Lamp5* GABAergic neurons).

ENVI performed well on ExSeq imputation and was only matched by gimVI in iterative leave-one-out validation, demonstrating that it can extend the ExSeq vocabulary beyond 42 genes (**Fig. 3c**). Its imputation of unimaged genes *Wfs1*, *Marcksl1* and *Pcp4* also matched Allen Brain Atlas ground truth (**Supplementary Fig. 5b**). *Marcksl1* spans multiple distinct cortical layers, highlighting ENVI’s ability to capture complex relationships between genes. ENVI imputation relies on the co-embedding of similar cell types in latent space for accurate inference. The stark difference in cell type frequencies between the ExSeq and matched scRNA-seq data poses a particular challenge, yet ENVI was able to accurately map single-cell and spatial samples to a unified embedding (**Fig. 3d**). To evaluate the latent space learned by ENVI, we assessed how accurately the embedding can transfer cell type label information between the scRNA-seq and ExSeq datasets. Since classical accuracy is biased towards more abundant populations when cell type frequencies vary, we quantified classification quality using balanced accuracy (BA), the average of the sensitivity and specificity of all classes. The ENVI latent achieved the highest BA, better reflecting cell type than the gimVI latent and more precisely transferring information than cell-type mapping by Tangram (**Fig. 3e**).

To provide a higher-throughput imaging benchmark, we tested ENVI on a primary motor cortex dataset generated with a 252-gene MERFISH panel, comprising 22 cell types from six imaged samples (18,515 cells) with matching scRNA-seq data (7,416 cells) from the Brain Initiative Cell Census Network^38^. Based on five-fold cross-validation, ENVI reconstruction of held-out genes more closely matched ground truth expression compared to Tangram, gimVI and NovoSpaRc (**Fig. 3f**), with strong statistical significance in most cases. We visualized cortical markers *Pardm8* and *Satb2* to investigate differences in imputation, and found that ENVI predicted expression correctly while Tangram only predicted expression in a subset of expressing cells and gimVI overestimated abundance in most cells (**Fig. 3g**).

### ENVI ascribes spatial developmental patterns to single-cell sequencing data

While the integration of spatial transcriptomics and scRNA-seq data makes it possible to infer the spatial patterns of thousands of genes, imputation can never replace direct measurement; moreover, highly multiplexed spatial technologies are within reach. A more valuable goal, then, is to add spatial context to the large number of scRNA-seq datasets being generated. In the same manner that we use the gene expression decoder to impute missing gene expression for the spatial data, we can use the spatial decoder in ENVI’s unique dual decoder scheme to infer COVET matrices for the scRNA-seq data, providing spatial context for data from dissociated cells (**Fig. 4a**).

**Figure 4.**
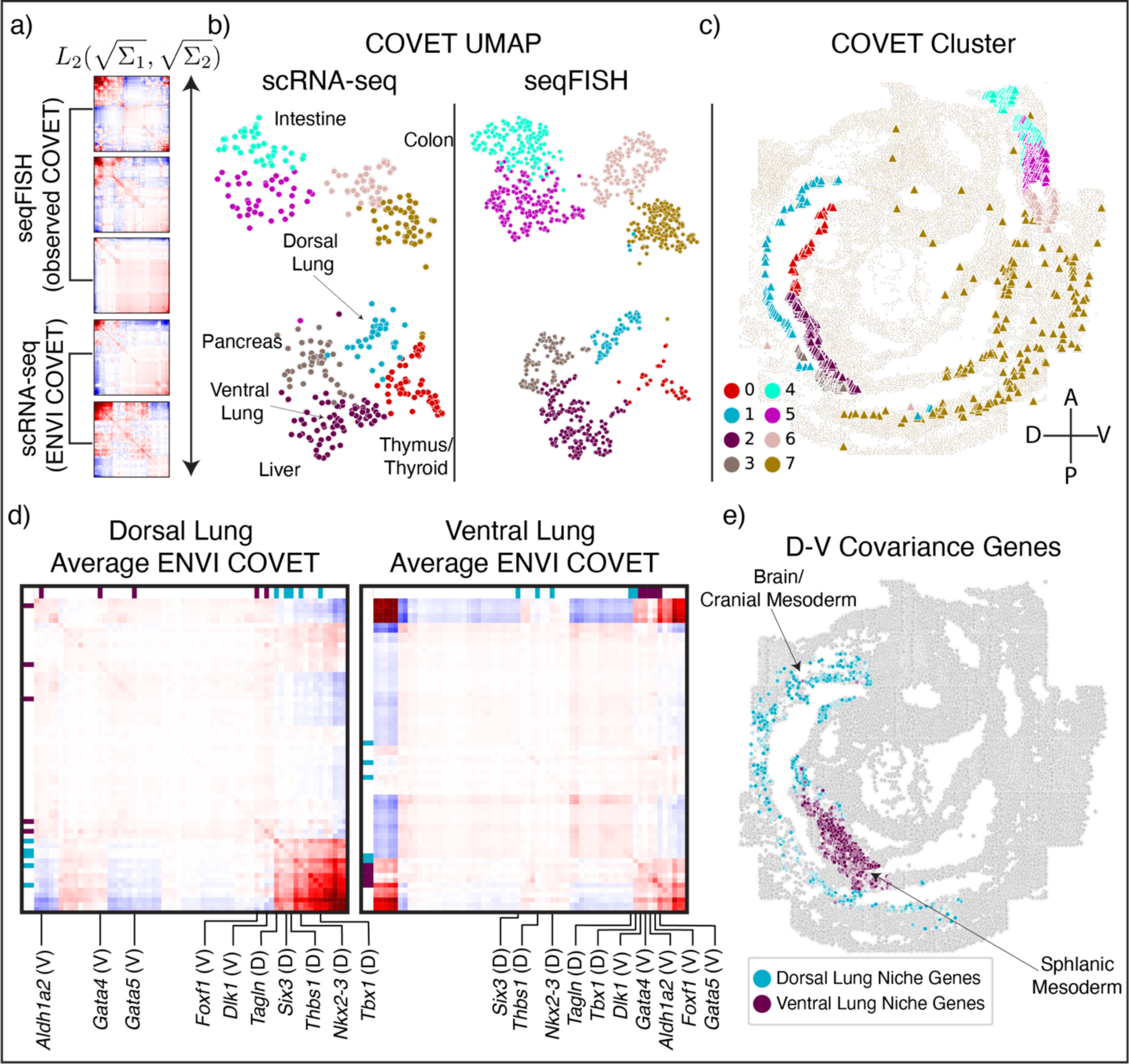
ENVI attributes spatial context to single-cell samples from mouse gut organogenesis. **a**, Schematic of COVET visualization using UMAP and clustering with PhenoGraph, calculated based on the AOT metric on both seqFISH and scRNA-seq modalities. **b**, UMAP of gut tube cells from scRNA-seq and seqFISH data, colored by PhenoGraph COVET cluster, showing coherence of cells with similar predicted neighborhoods. Developing organ labels in scRNA-seq data taken from Nowotchin et al.^48^ **c**, seqFISH gut tube data colored by COVET cluster, delineating spatial regions of the gut tube. **d**, Average ENVI-predicted COVET matrices for scRNA-seq data from dorsal lung (left) and ventral lung (right), sorted by hierarchical clustering. Gene clusters that are uniquely present in either dorsal or ventral lung are labeled. **e**, seqFISH cells near (but not part of) the gut tube, pseudo-colored by relative expression of unique dorsal or ventral lung genes highlighted by ENVI-predicted COVET matrices in **d**. Labels indicate predominant cell type in each colored region.

To test ENVI’s ability to predict spatial niches for single-cell data, we focused on gut tube cells within the mouse embryo dataset, which later give rise to the thymus, thyroid, lung, liver, pancreas, small intestine, and colon in a stereotypical anterior-to-posterior sequence. Although cells of the E8.75 gut tube are anatomically indistinguishable, organ-specific expression reveals that precursors are poised for their future fates, and that they are arranged in domains that prefigure organ emergence along the tube^48^. We asked whether ENVI can effectively capture this spatial information and project it onto dissociated scRNA-seq data.

To evaluate whether ENVI assigns the correct spatial context, we co-embedded ENVI-inferred COVET matrices for scRNA-seq data with observed COVET matrices for spatial data using the AOT metric, and clustered them^29^ into distinct niches containing cells from both modalities (**Fig. 4b**). It was difficult to assign organ primordia based on the very limited structure provided by seqFISH expression data alone (**Supplementary Fig. 6a**), but labeling scRNA-seq cells from the gut tube with organ-specific gene sets^48^ (Methods) revealed an almost one-to-one matching between organ precursors and COVET clusters (**Supplementary Fig. 6b**). We labeled the co-embedded seqFISH cells using the scRNA-seq-based organ annotations and found that the COVET clusters occupy distinct niches along the gut tube (**Fig. 4c**). Moreover, ENVI predicts the correct location of organ annotated COVET clusters from anterior to posterior; thymus and thyroid cells fall into the most anterior ventral region of the gut tube, followed by dorsal and ventral lung clusters, and finally, colon and small intestine.

Other integration methods do not model cellular environments, underscoring the difficulty of spatial inference. We tested gimVI and Tangram, which are dedicated to scRNA-seq and spatial integration, and Scanorama^49^, a general integration method commonly used for correcting batch effects in high throughput data, which relies on expression-based mutual nearest neighbors between two datasets. None could accurately label the seqFISH gut tube; gimVI failed to discern dorsal lung from thymus and thyroid, while Tangram and Scanorama labeling lacked any apparent spatial pattern (**Supplementary Fig. 6c**). ENVI was unique in assigning organ labels to seqFISH cells that matched their expected localization.

We next assessed whether ENVI-inferred niches can provide spatial information that goes beyond the endodermal cells of the gut tube. Focusing only on scRNA-seq data, we compared the inferred COVET matrices between ventral and dorsal lung COVET clusters and found distinct sets of genes that covary in each cluster’s average matrix—that is, genes that are coordinately expressed at high levels in their respective niches (**Fig. 4d**). *Dlk1*, *Gata4*, *Gata5*, *Aldh1a2* and *Foxf1* all covary in the ventral lung COVET cluster, but not the dorsal lung, whereas *Tagln*, *Six3*, *Thbs1*, *Nkx2-3* and *Tbx1* exhibit the inverse pattern. We plotted the average expression of these gene sets limiting to only non-endodermal cells in the seqFISH data, and found that ventral COVET genes are enriched in the ventral splanchnic mesoderm while dorsal COVET genes are expressed in the dorsal brain and cranial mesoderm (**Fig. 4e**). This observation validates the predicted DV spatial subdomains within the gut tube, and highlights the ability of ENVI to identify signaling niches, such as the mesodermal cells known to provide patterning cues to adjacent endodermal cells^50^.

### ENVI learns spatial gradients from single-cell data

While the spatial separation of the gut tube into primordial organs is relatively discrete, many developmental processes—such as the specification of spinal cord cells and their precursors, the neuroemesodermal progenitors (NMP), along the AP axis—are defined by continuous spatial gradients (**Fig. 5a**). Contrary to approaches based on discrete cell types, ENVI is quantitative and can capture these gradients. To highlight continuous spatial trends, we co-embedded empirical seqFISH COVET matrices with ENVI-inferred scRNA-seq COVET matrices for NMP and spine cells using a force-directed layout (FDL)^51^ and calculated their diffusion components (DCs) (**Fig. 5a**). The first DC is highly congruent with the AP axis (Pearson correlation 0.86), demonstrating that COVET can capture gradual trends in spatial data (**Fig. 5a,b** and **Supplementary Fig. 7a**). As the COVET DC is calculated from both seqFISH and scRNA-seq datasets, we can use it to assign AP pseudo-coordinates to NMP and spine cells from scRNA-seq data.

**Figure 5.**
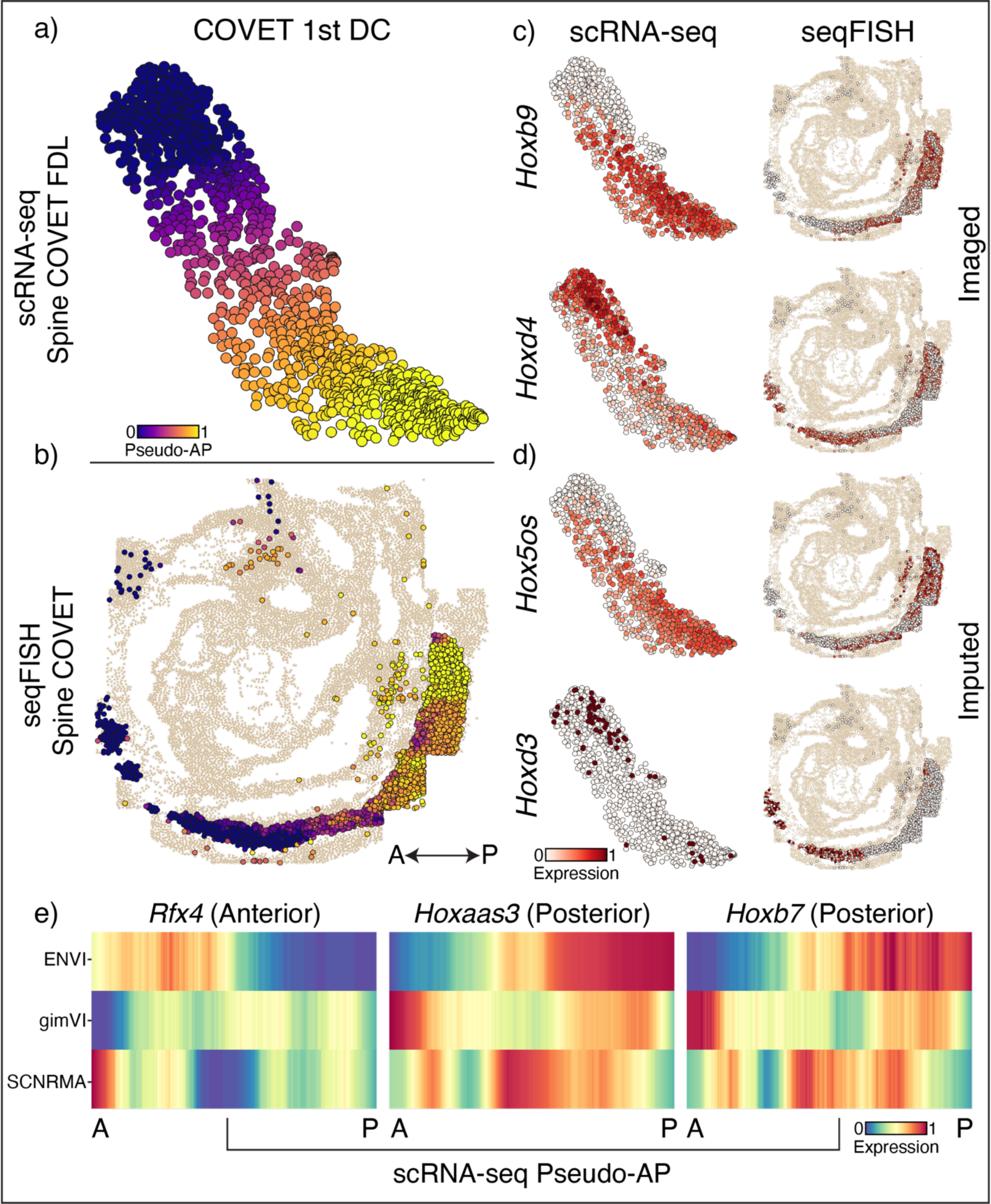
ENVI maps continuous spatial profiles of spinal gastrulation from development from single-cell and spatial data. **a**, Force-Directed Layout (FDL) based on ENVI-predicted COVET matrices of scRNA-seq spinal cord cells and their NMP precursors, colored by the first COVET diffusion component (DC). FDL and DCs were calculated using both the seqFISH and scRNA-seq COVET matrices. **b**, seqFISH NMP and spine cells colored by the first COVET DC, which traverses the AP axis. **c**, Expression of *Hoxd4* (anterior) and *Hoxb9* (posterior) markers in NMP and spine cells from scRNA-seq and seqFISH data, recapitulating expected AP localization. **d**, Same as **c**, colored by ENVI-imputed *Hoxb5os* (posterior) and *Hoxd3* (anterior) expression, as these markers were not imaged. **e**, scRNA-seq expression of canonical AP axis markers in NMP and spine cells ordered along pseudo-AP axis according to different integration methods. SCNRMA, Scanorama.

The first COVET DC reveals that scRNA-seq cells are correctly enriched for *Hoxd4*^48, 52^ (anterior) and *Hoxb9*^53^ (posterior) markers in their respective domains, consistent with seqFISH expression in NMP and spine cells (**Fig. 5c**). Furthermore, ENVI correctly mapped high expression of *Hoxd3*^52^ (anterior) and *Hoxb5os*^54^ (posterior) markers to scRNA-seq cells in the corresponding AP domains, demonstrating that ENVI modeling of the expression-environment relationship extends to genes that are not imaged (**Fig. 5d**). Conversely, ENVI-imputed *Hoxb5os* and *Hoxd3* expression for the seqFISH data mirrored the predicted spatial context of the scRNA-seq data.

We found that the major axis of variation between the COVET matrices that directly model the environmental niche reflect spatial organization of the tissue. By ordering NMP and spine cells along the first COVET DC, we recovered a pseudo-AP axis, which can be used to visualize predicted expression trends (**Fig. 5e**). Similar analysis of the gimVI latent space and Scanorama integration (Methods) led to inferior alignment with the true AP axis (**Supplementary Fig. 7b**), despite selecting the gimVI and Scanorama DCs most correlated (*r* = 0.76, 0.7 respectively) with true AP polarity (rather than the top component). This slightly lower correlation with the AP axis propagates into more pronounced inaccuracies in expression patterns; only ENVI correctly assigned *Rfx4*^55^, *Hoxaas3*^54^ and *Hoxb7*^56^ expression to the correct AP locations (**Fig. 5e**). More generally, canonical markers of both posterior and anterior regions are more correlated (or anti-correlated) with ENVI COVET pseudo-AP, compared to axes defined by gimVI and Scanorama (**Supplementary Fig. 7c**). ENVI can thus correctly uncover AP polarity within single-cell NMP and spine cells and correctly place them along this spatial axis.

### ENVI delineates tissue-scale spatial patterning in the motor cortex

Glutamatergic neurons are the most abundant cell type in the mouse MERFISH^38^ and corresponding scRNA-seq^57^ datasets. They are classified according to their distinct layer-specific localization and expression in the motor cortex, yet recent scRNA-seq studies have revealed greater heterogeneity within this broadly distributed class of neurons than previously appreciated^58^. We investigated whether ENVI can integrate data from multiple MERFISH samples and map them onto full transcriptome single-cell data to reveal tissue-scale insights and possible gradients in glutamatergic neuron patterning.

The MERFISH data includes examples of both discrete and continuous expression across the motor cortex, but it only consists of 252 genes. Despite the extreme rarity of some cell types in the scRNA-seq dataset, including only 19 oligodendrocyte precursor cells and 13 microglia, ENVI was able to harmonize both data modalities into a unified latent (**Fig. 6a**). We found that the first DC of ENVI-imputed COVET matrices is highly correlated with motor cortical depth, thus defining a ‘pseudodepth’ axis (**Fig. 6b**,**c**). Similar axes recovered from DC analysis of other integration methods were less accurate in recapitulating cortical depth (**Supplementary Fig. 8a–c**).

**Figure 6.**
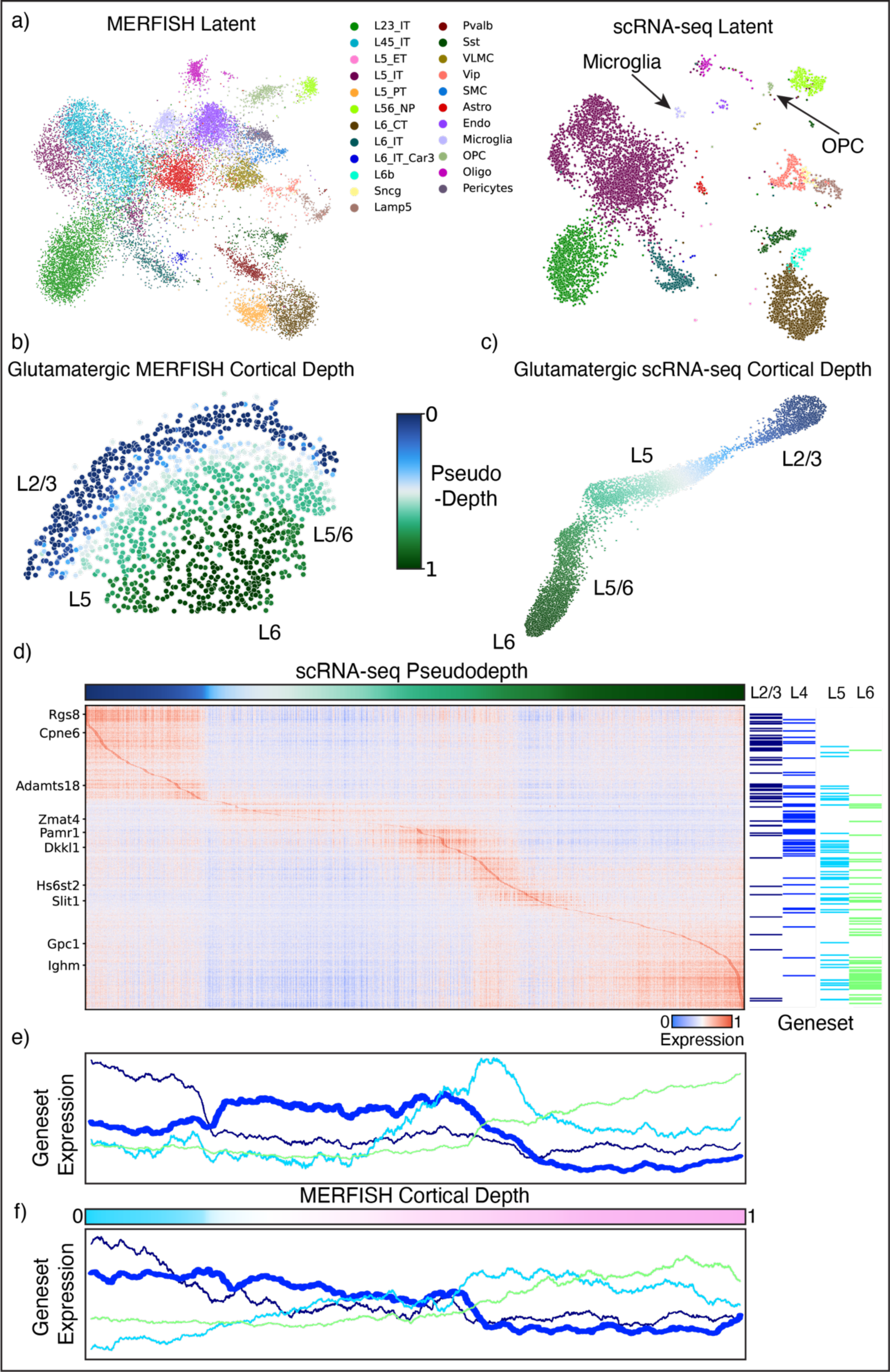
ENVI can extend gene expression gradients to the entire genome, across cortical tissues. **a**, ENVI latent embedding of motor cortex MERFISH (left) and scRNA-seq (right) data, labeled by cell types from Zhang et al.^38^ and Yao et al.^57^. OPC, oligodendrocyte precursor cells. **b**, Glutamatergic (excitatory) neuron from the MERFISH dataset, colored by the first COVET DC, representing pseudodepth. Cortical layers L2/3 to L6 are labeled. **c**, scRNA-seq COVET FDL of glutamatergic neurons, colored by first COVET DC. COVET FDL and DCs are calculated based on both scRNA-seq and MERFISH datasets. **d**, scRNA-seq expression of glutamatergic neurons, with cells (columns) and genes (rows) ordered by predicted cortical depth. Right, gene sets enriched in each cortical layer according to the Allen Brain Atlas. Labeled genes (left) are those for which ENVI-imputed expression matches ISH imaging in the primary motor cortex (**Supplementary Fig. 8d**). **e**, Average expression of cortical layer gene sets in each scRNA-seq glutamatergic neuron, ordered from left to right by predicted cortical depth, and colored as in **d**. L4 average expression is highlighted by a thick line. **f**, Average expression of the MERFISH panel markers present in each cortical layer gene set, for each MERFISH glutamatergic neuron. Neurons are ordered from left to right by true cortical depth.

Ordering cells by cortical depth and ordering genes by depth at peak expression provides a pseudospace heatmap of gene expression (**Fig. 6d**). The Allen Brain Atlas data (mouse.brain-map.org) validated the imputed COVET cortical depth of unimaged genes (**Supplementary Fig. 8d**), and mapping peak expression of Allen Brain Atlas gene sets to their imputed cortical depth more systematically validated ENVI’s inferred pseudospace.

Although the motor cortex is traditionally believed to lack L4 neurons, a recent study^59^ demonstrated a compartment between the L2/3 and L5 layers with the same basic synaptic circuit organization as L4 neurons in the sensory cortex. While the MERFISH data alone supports this, with a visible layer expressing L4 marker genes *Rspo1* and *Rorb*, our analysis provides much more extensive corroboration, predicting clear peaks for 64 L4 marker genes between the L2/3 and L5 layers (**Fig. 6d**). Moreover, analysis at the gene set level shows clear trends along predicted cortical depth (**Fig. 6e**), whereas the MERFISH gene panel is insufficient to resolve L4 signals from the other superficial layers (**Fig. 6f**). Thus, by mapping information about the environmental niche onto scRNA-seq data, ENVI can reveal spatial signals hidden in the partial transcriptome captured by multiplexed imaging data.

## DISCUSSION

ENVI robustly integrates scRNA-seq and spatial transcriptomics data, overcoming technical biases while retaining biological information. The algorithm provides superior performance for imputing missing gene expression in spatial modalities; it scales to millions of cells; and it has the distinctive ability to infer the spatial context of dissociated cells, even across multiple cell types in complex tissues. It thus uniquely provides a genome-wide view of spatial gradients across tissues.

ENVI’s capabilities critically rely on COVET as a representation of spatial niches. While most current spatial representations are based on cell typing and are thus discrete in nature, we developed COVET to take full advantage of the quantitative nature of gene expression measurements. The COVET matrix represents covariation between markers in a cell’s niche and uses optimal transport to derive a principled and quantitative model of cellular neighborhoods. The shift from discrete cell-type to continuous cell-state paradigms has been instrumental in single-cell genomics. COVET extends this notion to spatial data, enabling the discovery of continuous trends in microenvironments.

The performance of ENVI is primarily driven by three factors: 1) deep Bayesian inference to regress out modality-related confounders while learning nonlinear relationships between genes and niches, 2) explicit modeling of the entire transcriptome from scRNA-seq data, and 3) direct incorporation of spatial context via COVET. Whereas current methods only learn the common features (set of overlapping genes) between scRNA-seq and spatial datasets, the guiding principle of ENVI is to model all available information, and not rely on post-hoc inference from partial knowledge. This proves invaluable, as the ENVI model is imbued with both spatial context and full transcriptome information, allowing for reliable transfer of information between modalities.

Benchmarking on multiple different imaging technologies and tissue types shows that ENVI integrates data effectively, learning a latent space which is more reflective of true structure in the spatial and single-cell datasets, to successfully impute unimaged genes and reconstruct spatial context. The ENVI COVET space can correctly predict the niches of primordial organs from seqFISH and scRNA-seq data of mouse gastrulation, and COVET-based diffusion component analysis can highlight continuous anteroposterior trends of both expression and environment in the developing spine. By mapping scRNA-seq data from the motor cortex onto MERFISH cortical gradients, ENVI further establishes the existence of primary motor cortex L4 neurons, whose signal is hidden in the 252-gene MERFISH imaging data and requires full transcriptome data to uncover. ENVI-imputed scRNA-seq COVET matrices provide an accurate representation of both discrete and diffuse signals in the context of intact tissue, enabling spatial reasoning along the full transcriptome.

## CODE AVAILABILITY

ENVI and COVET are available as python packages at <u>github.com/dpeerlab/ENVI</u> and can be directly installed via pip with the command ‘pip install scENVI’. A Jupyter notebook with an ENVI tutorial which reproduces motor cortex MERFISH results is available at: <u>github.com/dpeerlab/ENVI/blob/main/MOp_MERFISH_tutorial.ipynb</u>.

## ACKNOWLEDGEMENTS

We would like to acknowledge Cassandra Burdziak and Esther Wershof for their insightful comments. This work was supported by NCI Cancer Center Support Grant P30 CA08748, NCI grant U54 CA209975, NCI grant R01 DK127821 (M.G.), NCI Human Tumor Atlas Network U2C CA233284 (D.P). D.P. is an HHMI investigator.

## AUTHOR CONTRIBUTIONS

D.H. and D.P. conceived the study and algorithm design. D.H implemented all algorithms used in this study and carried out testing, application, and data analysis. D.H., M.G. and A.-K.H. carried out data analysis and interpretation of the embryogenesis data. M.G collected the mouse embryogenesis HCR imaging data. D.H., T.N., and D.P. wrote the manuscript. D.P. supervised the study.

## METHODS

### Multiscale spectral similarity index

When comparing the spatial distribution of genes or markers across a tissue, it is imperative to have a robust metric that takes spatial structure into account. While ubiquitous metrics such a Pearson correlation, SSIM and root mean square error can provide some insight, they lack spatial context (e.g. cell-cell proximity or spatial patterns) and only measure per-cell discrepancy.

To devise a metric for spatial data, we borrow the multiscale structure similarity index measure (MS-SSIM)^42^, a ubiquitous metric for the quality of image reconstruction, from computer vision. Given two images, MS-SSIM iteratively downsamples each image, creating an image pyramid^60^, a multiscale signal representation consisting of the same image at multiple resolutions. MS-SSIM returns a weighted geometric average of the standard SSIM scores between the two images at each scale of the pyramid. Standard SSIM for two images, *x* and *y* is:

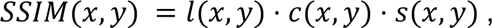

Where:

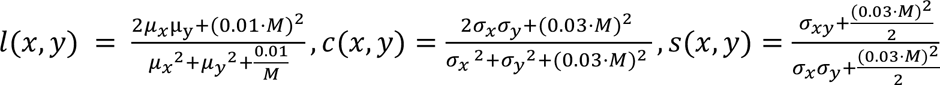

*M* represents the maximum values between *x* and *y*, and as and µ_x_ are their average values, µ_y_ and µ_x_ measure how each varies and µ_y_ represents how much they covary.

*l*(*x*, *y*), *c*(*x*, *y*) and *s*(*x*, *y*) are measures of ‘luminance’ (signal brightness), contrast, and structure, respectively. We note that while SSIM is meant for images, it can also be calculated between any two vectors of similar sizes.

We introduce the multiscale spectral similarity index (MSSI), an adaptation of MS-SSIM to spatial transcriptomics. The key difference is that we compare count matrices from segmented cells, rather than pixels, using a neighbor graph of spatially neighboring cells to capture structure. Intuitively, MSSI is a spectral analog of MS-SSIM; by rephrasing image coarsening to its graph-based counterpart, we can apply it to segmented cells and produce a multiscale, spatially driven score of expression reconstruction quality.

MSSI compares the expression profiles of two genes from a multiplexed image; 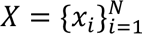 and 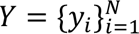 where the index *i* enumerates segmented cells and *x* and *y* can be either (1) two different genes or (2) a ground-truth gene and its imputed value. In addition, the spatial coordinate of each cell is 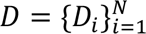. We first compute the k-nearest-neighbor graph *G*^1^ of segmented cells from 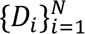. To generate a subsampled version of the kNN graph, we use a graph coarsening algorithm^61^, which pools nodes together based on their connectivity pattern, similar to how image downsampling groups pixels together (**Fig. 2c**). We iteratively coarsen and blur the graph four times by a factor of two, and produce the expression of each gene at each scale.

Mathematically, each coarsening step produces a pooled version of the graph 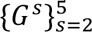 and a coarsening operator 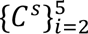, which is the mapping between nodes at one scale to nodes at the next and allows us to generate pooled versions of the gene expression signals:

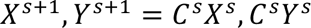

Following MS-SSIM, we compute the MSSI between the expression profiles at each scale and return their weighted geometric mean. In detail, we compute the *l*, *c* and *s* SSIM related values at at each scale, and derive MSSI based on their weighted product, as for the MS-SSIM:

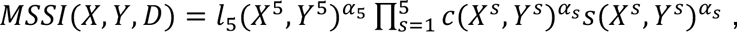

Where the weights are equivalent to those in MS-SSIM^42^:

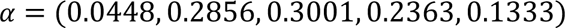

When *X*^i^, *Y*^i^ are anti-correlated (µ_xy_< 0), *s* is negative, which causes errors in computing the weighted geometric mean; we thus clip negative values to 0. This implies that if at any scale, *X*^s^, *Y*^s^ are anti-correlated, the MSSI will be 0, its lowest possible value. We also normalize the original-scale gene expression to be between 0 and 1, but we do not re- normalize at each coarsening scale.

### Spatial covariance representation

Our spatial covariance framework includes three components: the COVET statistic, similarity metric, and an algorithm to robustly and efficiently compute the COVET metric. The COVET framework assumes that the interplay between the cell and its environment creates covarying patterns of expression between the cell and its niche, which can be formulated via the gene-gene covariance matrix of niche cells. The COVET statistic constructs a shifted covariance matrix (which preserves algebraic properties of the covariance matrix) and thus enables the use of any measure of statistical divergence between covariances to define a principled quantitative similarity metric to compare niches. The key is to build the COVET statistic in such a manner that two COVET matrices are comparable, and to design a computationally efficient algorithm to quantify the statistical divergence between them.

The inputs to COVET are (i) the gene expression matrices (*X* ∈ *R*^6×8^), where *n* is the number of cells and *g* is the number of genes profiled, (ii) the location of each cell *in situ* and (iii) a parameter (*k*) that defines the number of nearest cells to be included in the niche. For each cell, we identify the *k* nearest cells based on their spatial proximity and construct a niche matrix 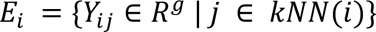 which represents the gene expression vector for each of those nearest neighbors. This produces an *n × k × g* tensor 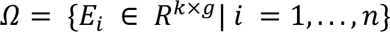, which are the niches of every cell. By default, we select *k* = 8, which usually captures the immediate niche of a cell, but the exact choice of *k* should reflect the dataset.

The fundamental goal of COVET is to transform those niche matrices into effective representations of a cell’s niche. To this end, we calculate the *shifted* gene-gene covariance matrix between cells in each niche matrix, where instead of using the classical formulation:

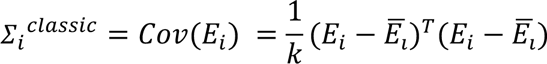

We swap the niche mean expression 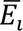 with the total expression mean 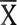(computing the mean over the entire dataset). This enables direct comparison between covariance matrices, as they are constructed relative to the same reference:

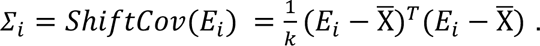

This creates a representation relative to the entire population, which can better highlight the features that are unique to each niche while also holding the same algebraic properties that the standard covariance matrix holds, namely being positive semi-definite. Therefore, we can harness measures of statistical divergence to derive a metric on the COVET matrices and quantify differences and similarities between niches. While we can conceptually use any statistical divergence measure, metrics like Kullback–Leibler (KL) divergence and Bhattacharyya^28^ distance are too computationally intensive and lack interpretability.

To meaningfully compare between niches, we cannot simply use the sum difference between two niche matrices *E*_i_ and *E*_j_,, as changing the cells’ order would change the result (whereas there is no meaning to any given order). An intuitive way to quantify niche similarity is by finding the best matching of cells between niche matrices by solving the assignment problem^62^. Optimal transport (OT)^63^ is a relaxed version of the assignment problem, where instead of matching cells one-to-one, OT finds the best *soft assignment* between cells. However, this approach to OT has no closed form solution and does not scale to large datasets. Instead, we can use the closed form solution of OT between covariance matrices, known as the Fréchet distance^27^:

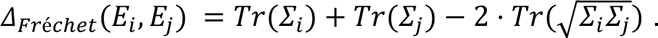

The Fréchet distance has time complexity of *O(k^3^)*, which would require billions of these pairwise computations between all niches for the analysis of large-scale datasets and is thus computationally intractable. To further speed up computation, we swap the matrix square-root (MSQR) and product operation in the last term of the Fréchet distance, and define the *approximate optimal transport* (AOT) distance as:

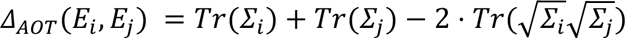

If Σ_i_ and Σ_j_ are commutative, this is no longer an approximation and 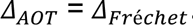. While in the form above, both the approximate and true Fréchet distance require O(*k*^3^) operations between each pair of niches, and O(*k*^2^) to compute the full distance matrix, using the fact that *Tr*(AB) = ||A ⊙ B||, where ⊙ marks element wise multiplication, we arrive at:

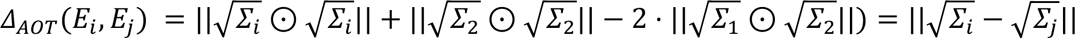

Therefore, when working in square-root space, we do not require any computationally extraneous matrix multiplication and many calculations of MSQR. Instead, we first calculate the MSQR of each COVET matrix, which is O(n*k*^3-^), and then simply calculate pairwise Euclidean distance, for a total time complexity of O(n*k*^3^+n^2^*k*^2^), which is significantly more efficient than O(*k*^2^) for large n.

Since AOT can be formalized as the *L*_2_ between MSQR of COVET matrices, it allows for direct use of any algorithm which is based on the Euclidean distance, such as UMAP, tSNE^64^ and FDL^51^, clustering^29^, and diffusion component^30^ (DC) analysis. We can simply compute MSQR of the COVET matrices, flatten the resulting matrices into 1D vectors, and apply the default implementations of all the mentioned algorithms. We can further leverage the Euclidean distance representation of the AOT metric and use computational accelerators designed to compute classical pairwise distances for additional speed gains.

We demonstrate that AOT is a good approximation by benchmarking against the true Fréchet distance and the Bhattacharyya distance, another common metric for distances between covariance matrices. Across various sizes of random sets of 64 by 64 covariance matrices, we test the runtime to compute the 10 nearest-neighbors matrix in covariance space. As covariance matrices are positive semi-definite, to randomly generate *n* covariance matrices of 64 by 64 elements, we first sample *n* random 64 by 64 matrices (using the standard normal), and multiply each by its transpose, as a matrix Gramian is always positive semi-definite:

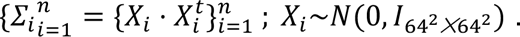

We find that while AOT produces accurate similarities, its runtime is at least an order of magnitude smaller than that of other metrics, with Fréchet and Bhattacharyya failing on sample sizes larger than 3,000 matrices due to out-of-memory error. Using a GPU implementation of kNN distance built for the Euclidean metric, which can be easily adapted for AOT, the spatial covariance metric is indeed scalable to massive datasets, taking under a minute to compute the kNN matrix between 100,000 samples (**Supplementary Fig. 1a**).

On real COVET matrices, calculated from the eight nearest neighbors of the splanchnic mesoderm cells from the SeqFISH assay^35^, using the 64 most highly variable genes from the total 350 imaged genes, we find high concurrence between the three metrics. We calculate the pairwise distance between the splanchnic mesoderm COVET matrices according to each metric which we use to calculate UMAP 2D embeddings and PhenoGraph clusters. We find high qualitative congruence between AOT, Fréchet and Bhattacharyya, indicating that AOT does not lose quality to gain computational efficiency (**Supplementary Fig. 1b,c**).

### ENVI algorithm

The ENVI (ENvironmental Variational Inference) algorithm integrates scRNA-seq and spatial transcriptomics data in a manner that can infer spatial context for scRNA-seq and missing genes for spatial transcriptomics. The core assumption of ENVI is that the COVET matrix, which captures the interplay between a cell’s phenotype and its microenvironment, provides information that better empowers the data integration, captured with multiplexed RNA imaging technologies, and encapsulated by COVET.

ENVI is grounded on autoencoder variational inference, but diverges from previous work^7, 23^; instead of modeling only common features between modalities, it explicitly models both the microenvironment context for spatial data and the full transcriptome expression of scRNA-seq. Unlike prior work, ENVI trains the variational autoencoder (VAE)^25^ to both reconstruct full transcriptome expression and spatial context from partial transcriptome samples.

To integrate between scRNA-seq and spatial data, ENVI learns a common latent space for both data modalities by marginalizing the technology-specific effect on expression via a conditional variational autoencoder (CVAE)^26^. It achieves this by augmenting the standard VAE with an auxiliary binary neuron in the input layers to the encoding and decoding networks representing each data modality. Integration is crucial, as there are fundamental technology-specific artifacts in each modality (**Supplementary Fig. 2a**). To learn the combined latent space, ENVI takes as input the scRNA-seq count matrix *X*_3;_ with n_3;_ cells and their full transcriptome of *g_sc_* genes, and the segmented expression profiles *X*_3E_ from n_3E_ cells and *g_st_* imaged genes. ENVI then uses the spatial data to compute the COVET matrix for each cell and their MSQR to align with the AOT distance formulation.

Our next step is to use a conditional autoencoder to build a shared latent space for both data modalities. As the combined embedding must incorporate spatial context and full transcriptome information, and must remove confounders relating to modality, we set the latent dimension to 512, substantially larger than standard VAEs in single-cell genomics, which usually have around ten neurons^23, 32, 33^. As input to the encoder, ENVI takes either spatial or scRNA-seq samples (the latter reduced to the subset of genes that have been imaged), along with the auxiliary neuron *c* having value *0* for the spatial data and *1* for scRNA-seq. The expression profile along with the auxiliary neuron are transformed into the latent variable *l* using the same encoding neural network, regardless of data modality:

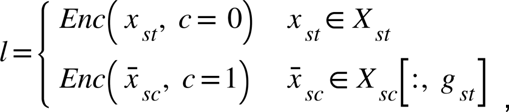

where the encoder returns two vectors µ_*l*_and σ_l_, which parameterizes a Gaussian with diagonal covariance describing the posterior distribution of the latent. To calculate gradients through random samples, we utilize the reparameterization trick, which involves generating a sample from the standard normal ε∼N(0,1) and describing the latent through a function of ε,µ_1_ and σ_1_and treating ε as a constant:

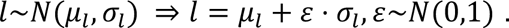

Through the training process, our goal is to have the latent encode not only gene expression, but also information about the spatial context of a given cell, while removing confounding effects to allow transfer learning between modalities. This is achieved by optimizing a single latent space to accurately decode both the full transcriptome and COVET matrix for both data modalities, each missing one of these components. The requisite that the latent space be capable of decoding the spatial niche is a novel aspect of ENVI and imbues sufficient spatial information into the latent space during training.

The latent of either modality, along with the appropriate auxiliary neurons, is fed into the *expression* decoder network *Dec_Exp_*. The loss function, calculated by comparing the activations in the output layer to the true expression profiles, needs to reflect the underlying distribution of each data modality. As scRNA-seq suffers from overdispersion and dropout, we use the negative binomial distribution to model the data, similar to previous work^23, 34^. During training, the scRNA-seq data provides transcriptome-wide expression; therefore, we can include genes whose expression was not provided to the encoder in the loss function, allowing our encoder to model genome-wide expression.

The negative binomial distribution has two parameters per gene—the number of failures, *r*, and success probability, *p*. Thus, the output layer of the decoder consists of 2 ⋅ *g*_sc;_ neurons, where the first *g*_sc;_ neurons are the parameter *r* and the latter *g*_sc;_ are *p*, using the *so*ftplus nonlinearity for *r* and the sigmoid function for *p* to keep it a valid probability:

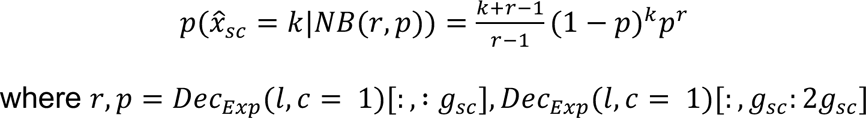

Multiplexed imaging based on FISH tends to have a high molecular capture rate^3^. Therefore, we use the Poisson distribution to model spatial data, and have the first *g*_st_ neurons in the output layer parameterize the per-gene rate parameter λ using s*o*ftplus nonlinearity to ensure it is a valid rate value:

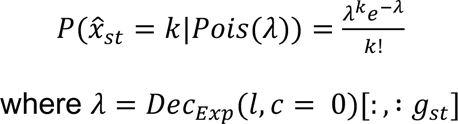

A standard CVAE, in which all neural parameters are shared aside from the auxiliary neurons, is sufficient to simply integrate between scRNA-seq batches, as demonstrated in scArches^32^. However, to successfully integrate scRNA-seq and multiplexed FISH-based technologies, a single auxiliary neuron is not sufficient to reasonably regress out all biases. In ENVI, only the first *g*_3E_ neurons of the output layer are shared by the two data modalities, with the rest being solely trained on the scRNA-seq data. Those additional technology-specific parameters improve the ability of ENVI to regress out confounders from the latent embedding, beyond the auxiliary neuron.

Finally, to reconstruct the spatial COVET matrices, we include an additional *environment* decoder network *Dec*_Exp_ whose role is to reconstruct the COVET from the latent, which can be trained from the spatial data. The output layer of the environment decoder has 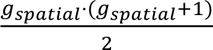 output neurons parameterizing the lower triangular Cholesky factor. The Gramian matrix of the output layer is the mean parameter of a standard normal, reflecting our AOT distance, as the log-likelihood of the standard normal is the *L*_2_ distance.

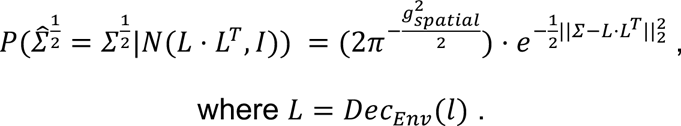

The output of the environment decoder is the MSQR of the COVET matrix, which is trained to minimize the *L*_!_ error with the MSQR of the true COVET matrix. Naively using the AOT metric in this manner involves computing the MSQR of the COVET samples during training, which can be computationally prohibitive. Instead, we first calculate the MSQR of all COVET matrices, which ENVI is directly trained to reconstruct.

We train ENVI simultaneously on samples from both spatial and single-cell datasets, using mini-batch gradient descent on the variational inference loss. With the learned ENVI model, we impute missing genes for the spatial data by treating the latent embedding of the spatial data as if it were from the single-cell data, using the single-cell auxiliary variable and parameterizing as a negative binomial instead of a Poisson. Conversely, we reconstruct spatial context for the single-cell data by applying the *environment* decoder on its latent, as if it was the latent of the spatial data.

In more detail, we train ENVI to optimize the evidence lower bound (ELBO) with a standard normal prior on the latent, with the goal of increasing the likelihood of the observed data {*X*_sc_ *X*_st_ Σ_st_} for the parameterization of their decoded distributions 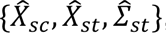, while minimizing the KL divergence between the latent distribution and *N*(0,1):

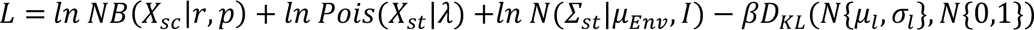

To train ENVI to impute missing genes for the spatial data, we generate the latent embedding *l*_st_ by passing *X*_st_ through the encoder, and run the latent layer through the *expression* decoder, but with the inverse auxiliary neuron, as if the embedding came from scRNA-seq data:

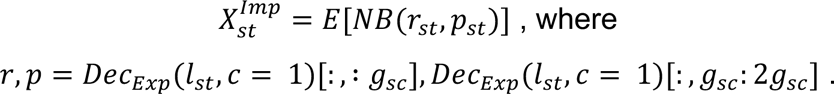

Similarly, we reconstruct the spatial context for dissociated scRNA-seq samples by passing the scRNA-seq latent embedding *l*_sc;_ through the *environment* decoder:

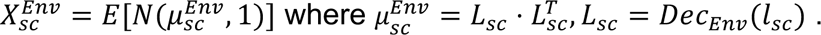

We have implemented multiple distributions in ENVI, including normal, Poisson, negative binomial and zero-inflated negative binomial (ZINB)^65^, which can be chosen for either modality to reflect preprocessing steps or varying levels of noise or dropout. The rate or mean parameters (λ for Poisson, *r* for NB and ZINB and µ for normal) must all be defined per-cell and per-gene, and shared across the single-cell and spatial data. However, all other parameters can be chosen to be either per-cell and per-gene, or simply per-gene, and can be shared between technologies or distinct.

By default, the encoder and two decoder networks consist of 3 hidden layers, each with 1,024 neurons and the latent variable consists of 512 neurons. The latent embedding consists of 512 neurons and the prior coefficient is set to β = 0.3. We train ENVI for 2^14^ gradient descent steps with the ADAM optimizer^66^ with learning rate 10^-3^ (lowered to 10^-4^ during the last quarter of training steps) and a batch consisting of 1,024 samples, half coming from scRNA-seq and the other half coming from the spatial data. To reduce computational complexity, we subset the scRNA-seq dataset to the union of the 2,048 highly variable genes and all genes included in the spatial dataset, rather than the full transcriptome.

Training ENVI takes on average less than 10 min of a single 12 GB GeForce RTX 2080 GPU, and less than 30 min without a GPU. The training of ENVI is constant in both time and memory regardless of dataset size. By contrast, methods such as Tangram and NovoSpaRc scale quadratically with dataset size and cannot be accelerated with GPU on datasets larger than approximately two thousand cells.

We benchmarked the run times of ENVI, Tangram^7^, NovoSpaRc^24^ and gimVI on spatial and scRNA-seq datasets of various sizes. Datasets were randomly generated, sampling from the Poisson distribution with parameter λ = 10 to create an *m* cells by 128 genes spatial dataset and from a distribution with parameters *r* = 10, *p* = 0.8 for a *m* cells by 2048 genes scRNA-seq data, where *m* is the parameter we benchmark against. Genes were named by their order, so that the first 128 features in the random scRNA-seq dataset were the genes in the spatial data. All models were trained with their default parameters, except that Tangram was trained on a CPU when *m* was larger than 10,000 cells, as it produced an out-of-memory error on our standard 12GB GPU. Model training was stopped prematurely if it exceeded 5 h.

As expected, ENVI’s training time was consistently around 10 min, regardless of dataset size (**Supplementary Fig. 2b**). The run time of gimVI grew linearly with dataset size as expected, since gimVI is trained for 200 epochs over the datasets. Unlike ENVI, which relies on deep variational inference and is not directly impacted by dataset size, NovoSpaRc and Tangram learn a cell-to-cell mapping between the spatial and single-cell datasets. While somewhat efficient on small datasets, we found that NovoSpaRc was very slow on large samples. Similarly, on larger spatial and scRNA-seq datasets, we found that GPU acceleration is not possible for Tangram, and as for NovoSpaRc, training was prohibitively slow.

### Whole-mount hybridization chain reaction

Whole-mount HCR mRNA in situs were performed as described previously^47^, with minor modifications^67^. Midgestation embryos at embryonic day (E) 8.75 were treated with 10 µg/ml proteinase K for 5 min at room temperature followed by washing and post-fixation in 4% PFA for 20 min. Embryos were incubated in hybridization buffer supplemented with 2 pmol of each probe (*Ripply3*, *Nkx2.5* or *Tlx2*) overnight at 37°C, followed by an amplification step with 60 pmol of each fluorophore-conjugated hairpin for 12-16 h at room temperature. Embryos were then stained with 0.5 µg/ml DAPI (Thermo Fisher Scientific) and cleared using a modified Ce3D^+^ clearing protocol^68^ for 24–48 h. Images were acquired on a Nikon A1R laser-scanning confocal microscope with a 10x objective and 3.0 µm z-step size. Image rendering and optical sections were generated using IMARIS (v9.9.0, BitPlane). All probes, hairpins and buffers were designed and purchased from Molecular Instruments, Inc.

### Benchmarking imputation

To benchmark ENVI gene imputation against alternative methods, we follow a similar scheme to previous approaches^23, 41^, where a test set of held-out genes is generated using cross-validation, and imputation of held-out genes is compared to true expression. For imputation metrics, we opt to use both the widely used Pearson correlation and our spatially aware multiscale spectral similarity index (MSSI) metric. We evaluate log expression and imputation profiles, with pseudocount 0.1.

While many algorithms now use scRNA-seq data to predict the expression of missing genes in spatial transcriptomics datasets^39, 40, 69–71^, we compared ENVI to gimVI and Tangram for their competitive performance in a recent benchmarking study^41^, and NovoSpaRc for its use of spatial context and optimal transport when integrating spatial and single-cell data.

For imaging datasets with relatively few genes, such as osmFISH and ExSeq (limited to a few dozen genes), we perform a full leave-one-out cross-validation experiment. One by one, each gene in the imaging panel is hidden, and its expression is predicted by each algorithm. For datasets from higher-throughput imaging modalities such as seqFISH and MERFISH, which assay hundreds of genes, we opt for five-fold cross validation. The imaged gene set is randomly divided into five groups, and each model is trained on 4 groups and tested on the withheld group. Finally, to appraise which model is superior for each dataset, we use a *relative* one-sided t-test, as scores are paired across genes.

We benchmark all models using their default parameters and instructions on all datasets:

- gimVI: Per the scvi-tools webpage (docs.scvi-tools.org), we train gimVI for 200 epochs with a batch size of 128 and latent dimension. The spatial and scRNA-seq datasets are parameterized with the NB and ZINB distributions, respectively. To impute genes with the trained model, we follow the instructions in the manuscript and train a kNN regression model on the scRNA-seq latent and full transcriptome expression, with *k* being 5% of the number of cells in the single-cell dataset. We then apply the regression model on the spatial data latent to predict the expression of the unimaged genes.
- Tangram: We use default parameters (https://github.com/broadinstitute/Tangram) and train the model for 1,000 epochs using the ‘cells’ mode. We set the density prior to be uniform, as all spatial datasets we benchmark against are single-cell resolution. With the learned mapping, we use the tangram *project_genes* function to impute genes from the scRNA-seq onto the spatial dataset.
- NovoSpaRc: We follow the repository instructions (https://github.com/rajewsky-lab/novosparc), and use an alpha coefficient on a spatial location prior of 0.25 and smoothness parameters *epsilon* of 0.005. To compute the scRNA-seq pairwise distance matrix, we subset the dataset to the union of the 2,048 most highly variable genes and all the genes included in the spatial dataset. For spatial datasets consisting of multiple different samples, such as the MERFISH motor cortex which has 6 distinct slices, we train a different model on each sample, as NovoSpaRc does not handle multiple samples. To impute missing genes, we use the *tissue.sdge* function, which applies the learned mapping.

### Application of ENVI to SeqFISH embryogenesis dataset

We showcase ENVI on the embryogenesis seqFISH and scRNA-seq datasets, with the goal of demonstrating how ENVI can be applied to tissues with complex spatial patterns. In the gastrulation data^35^, the authors used seqFISH to image 351 genes in tissue sections of mouse embryos from embryonic day 8.75 (E8.75), resulting in 57,536 cells imaged. This data was paired with scRNA-seq data collected from mouse at E8.75 consisting of 12,995 cells^36^. Starting with the originally published preprocessing, we further processed the scRNA-seq data by removing mitochondrial genes and genes expressed in under 1% of cells. In addition, we excluded cells with library size greater than 33,000 (set manually to match the knee point), along with cells annotated as ‘nan’ or representing doublets. To avoid any confounding batch effects^72^ in the scRNA-seq dataset, we only use the largest sequencing batch (labeled as ‘3’). For the seqFISH dataset, we only used the first (‘embryo1’) of three imaged embryos, removed cells with abnormally high total expression (threshold set manually to 600) and removed the gene *Cavin3*, which did not appear in the scRNA-seq dataset. For both datasets, we used the cell-type annotations provided by the authors and visualized the seqFISH data using the spatial coordinates and scRNA-seq dataset using a UMAP embedding (**Fig. 2a**).

To apply ENVI, we took the union of the 2,048 most highly variable genes in the scRNA-seq data, the seqFISH measured genes, all HOX genes, and several organ markers (**Supplementary Table 1)** and trained the model with default parameters. We visualized the learned latent posterior of the seqFISH and scRNA-seq datasets using UMAP and found that cell types tend to co-embed regardless of modality (**Fig. 2b**). Incongruences in the latent mostly stemmed from differences in nomenclature between seqFISH and scRNA-seq labelings (e.g., paraxial mesoderm in the single-cell dataset is labeled cranial mesoderm in the spatial; somitic and presomitic mesoderm refer to the same tissue).

To test the imputation of non-imaged canonical organ markers *Ripply3*^44^ (lung), *Nkx2-5*^45^ (heart) and intestine *Tlx2*^46^ (intestine), we visualized their imputed z-scored, logged expression and thresholded values lower than 2, finding these to be expressed almost exclusively in the correct organ (**Fig. 2f**). To confirm that these genes are expressed in the correct location at E8.75, prior to organ formation, we imaged each organ marker gene on separate embryos using whole mount HCR (**Fig. 2f**, see section Whole-mount hybridization chain reaction). HCR produces per-gene 3D images, which we orient coronally to match the seqFISH data. We similarly trained gimVI and Tangram on the complete scRNA-seq and seqFISH datasets to impute *Ripply3, Nkx2-5* and *Tlx2*. Visualized in the same manner as our ENVI imputation, we compared which imputation best matches the expected organ expression. For each gene, ENVI imputation more closely matches the experimental data (**Supplementary Fig. 3b**).

### osmFISH imaging of somatosensory cortex

We used a 33-gene osmFISH imaging dataset of the somatosensory cortex, which includes a single osmFISH image comprising 4,530 cells and 3,005 cells profiled with scRNA-seq^3^, utilizing preprocessing and cell type annotation provided by the authors (**Supplementary Fig. 4a**). We refrained from any additional processing of the data besides removing genes that appeared in less than 1% of cells in the scRNA-seq data. As the osmFISH data was more dispersed than the MERFISH and seqFISH datasets (**Supplementary Fig. 2a**), we opted to model the spatial data with the negative binomial (NB) distribution instead of the Poisson. Due to the limited size of the scRNA-seq dataset, we changed its parameterizing distribution from NB to ZINB. Since the total sample size is small (combined datasets have fewer than 10,000 cells), we also increased the reliance on the prior latent distribution and increased the regularization to β = 1.0, which is common practice in Bayesian modeling.

We compared ENVI and gimVI learned latent spaces based on annotated cell types from the osmFISH and scRNA-seq datasets. We visualized the latents of each with a 2D UMAP embedding. The ENVI embedding separates distinct cell types, with similar labels from the two data modalities occupying similar spaces (**Supplementary Fig. 4b**), whereas gimVI confuses oligodendrocytes and pyramidal neurons and cannot accurately co-embed osmFISH and scRNA-seq endothelial cells (**Supplementary Fig. 4c**). We quantified integration quality and calculated the average center-of-mass embedding for each cell-type, from both seqFISH and MERFISH datasets, in the gimVI and ENVI embedding spaces. ENVI and gimVI latent dimensions are vastly different in size (512 for ENVI compared to only 10 for gimVI), so for a fair comparison, we normalized each column in the pairwise distance to a maximum value of 1. In the ENVI latent, the center of mass for each osmFISH cell type is distinctly closer to its counterpart in the scRNA-seq data compared to other cell types, whereas cell types are less well separated in the gimVI latent (**Supplementary Fig. 3d**). For each cell in the scRNA-seq data, we quantified this as the ratio of its 5 osmFISH nearest-neighbors in the latent space that share its cell type, and averaged across the 6 cell types. The latent cell type agreement is 0.58 for ENVI and 0.38 for gimVI.

We further imputed the expression of 3 unimaged genes onto the osmFISH dataset using the full ENVI model (**Fig. 3b**), and validated by comparing the (logged) expression profiles to the Allen Brain Atlas ISH images of the somatosensory cortex (**Fig. 3b**). ENVI imputation and Allen ISH images both specify *Dti4l*, *Rprm* and *Ndst* expression in the L2/3, L5 to L6, and CA1 regions, respectively. The Allen atlas provides both raw ISH images and processed, cell segmented expression profiles. Since we found each view difficult to interpret on its own, we overlaid the processed profiles on top of the raw ISH images for clarity.

### ExSeq of visual cortex

We selected a visual cortex dataset profiled by ExSeq^37^ as an example of in situ sequencing instead of ISH. ExSeq can image tens of genes at nanometric resolution in a single tissue. ExSeq was used to image 42 genes capturing 1,271 cells, with a paired scRNA-seq dataset comprising 13,586 cells^37^. We used the preprocessing in the original publication, and in the scRNAseq data further filtered genes appearing in less than 5% of cells and normalized the total library size of each cell to 10,000. Both the spatial and single-cell datasets were labeled into distinct neuronal and non-neuronal compartments based on cell-type labeling by the authors. The spatial data was visualized by in-situ coordinates and single-cell expression profiles by 2D UMAP embeddings (**Supplementary Fig. 5a**). We benchmarked performance on the ExSeq dataset using leave-one-out cross validation. We also trained ENVI on the complete ExSeq and scRNA-seq datasets, and imputed the expression of the cortical layer genes *Wfs1* (L2/3), *Marcksl1* (L5), and *Pcp4* (L6) (**Supplementary Fig. 5b**).

To quantify the quality of ENVI integration of spatial and scRNA-seq data, we visualized and compared each embedding with UMAP (**Fig. 3d**). Since both modalities come with ground-truth labels, we can directly measure the accuracy of transferring cell type labels from the scRNA-seq to the ExSeq dataset. We benchmarked against gimVI, whose learned latent can also be used to transfer knowledge from single-cell to spatial and vice-versa, and Tangram, whose trained cell-cell mapping is designed to label spatial data. Since NovoSpaRc does not assign cell type labels, it was not included in the comparison:

- ENVI: To transfer labels from scRNA-seq onto the ExSeq data, we fit a kNN (*k* = 5) classifier to predict scRNA-seq cell types from the ENVI latent, and use it to assign labels to the ExSeq data from its ENVI latent.
- gimVI: We train gimVI with the default parameters laid out in *Benchmarking Imputation*, and apply the exact same latent kNN based procedure we used to label the ExSeq cells with ENVI, but instead using the gimVI latent.
- Tangram: We fit Tangram (default parameters, see *Benchmarking Imputation*), to learn a mapping between the scRNA-seq to the ExSeq data. Unlike gimVI and ENVI, Tangram does not provide a unified embedding, but instead transfers information using its latent mapping. We apply Tangram’s *project_cell_annotations* function which assigns each cell type a probability of being mapped to each ExSeq cell according to the scRNA-seq data. We label each ExSeq according to the most probable cell type mapped to it.

We compare the predicted cell types to their ground-truths using the balanced accuracy, which is more suitable for unbalanced classification since cell type proportions are not uniform. We find that ENVI reproduces the original cell types better than both gimVI and Tangram, showing that its integration of the ExSeq and scRNA-seq datasets is more reliable (**Fig. 3e**)

### Spatial organization of emerging organs

By applying the trained COVET spatial decoder on the latent encoded from scRNA-seq data, ENVI infers its spatial context. We tested this by focusing on endodermal (gut tube) cells from the embryogenesis dataset (see above for how ENVI was trained on this data). At E8.75, endodermal organs are not yet formed, but cells from scRNA-seq data already cluster into primordial organs, ordered according to the location from which they will emerge along the gut tube^48^. Using a deeply profiled endoderm scRNA-seq atlas as reference^48^, we find gene sets associated with each organ using the *rank_genes* function implemented by *scanpy*^73^, which uses a Wilcoxon test to find differentially expressed genes in each organ (**Supplementary Table 2**). As the thymus and thyroid are not well delineated at this stage, we collapsed the two labels in the reference to thymus/thyroid.

We used those gene sets to classify the gut tube cells from the co-embedded scRNA- seq^36^ into 7 emerging organs: thymus/thyroid, dorsal lung, ventral lung, liver, pancreas, colon and small intestine. We co-embedded the ENVI imputed COVET matrices from the scRNA-seq data and the measured COVET matrices from the seqFISH data with UMAP using the AOT distance. We then used PhenoGraph to cluster the dataset into 12 clusters and labeled each cluster with its best matching organ based on z-scored and logged expression of each gene set, averaged across all cells in that cluster. Most clusters were highly distinct, while some co-expressed several programs. Clusters for which the (z-scored) ratio between the highest and second-highest-expressed gene set was more than 1.5, were labeled with the most highly expressed organ. To decide ambiguous clusters with ratios lower than 1.5, namely C5, C6 and C7, we inspected marker gene expression (*Ripply3*^44^ for lung, *Pdx1*^74^ for pancreas) individually and assigned a label among the most expressed gene sets.

- C5: The highest expressed organ in C5 was thymus/thyroid, but since the average expression of *Ripply3* and *Irx1*^75^ (lung markers) was high (average z-score logged expression was 0.90) while that of *Nkx2-1*^45, 76^ (thymus/thyroid marker) was low (- 0.15), we labeled the cells of C5 as ‘dorsal lung’ (the second-highest expressing organ).
- C6: The most expressed gene set was dorsal lung, with pancreas being a close second. Since the cluster had minimal expression of *Ripply3* and *Irx1* (0.18), while being enriched for *Pdx1* (0.43), C6 cells were assigned to the pancreas.
- C7: The first and second most highly expressed organs were pancreas and liver, respectively. Due to high *Pdx1* expression (0.99), and low expression (-0.12) of liver marker *Ppy*^77^, we kept the pancreas label for this cluster.

We quantified the agreement between reference and query labeling by calculating the pairwise Pearson correlation between the average (z-scored and logged) expression of all genes in the organ gene sets for each dataset. Across all organs, the mean correlation between query and reference expression was 0.74, whereas the background average pairwise correlation between any two organs across the two datasets was 0.006.

Using a similar approach to classify endodermal cells from the seqFISH data proved unsuccessful. For each gene set, we keep only the genes that were included in the seqFISH panel and visualized their average (logged and z-scored) expression on a 2D UMAP embedding of the seqFISH gut tube cells (**Supplementary Fig. 6a**). While there is a separation between the anterior organs (thymus/thyroid, lungs, liver and pancreas) and the posterior organs (small intestine and colon), there is too much overlap in expression within organs from each compartment to reliably assign identities. We attempted to label seqFISH gut tube cells via PhenoGraph clusters and assign organs based on the average expression of each gene set (subset to seqFISH assay genes only), but almost all seqFISH clusters had ambiguous expression and the ratio between the highest and second-highest expressed gene set was lower than 1.5, so we simply assigned organs based on the most highly expressed genes. The average Pearson correlation between the average expression of each cell type was 0.55 (average pairwise was 0.007), distinctly lower than the average correlation with the scRNA-seq labeling (0.74).

As an additional comparison between the seqFISH and scRNA-seq organ assignments, we calculated the assignment *significance*, which we define as the ratio between the correlation of the average expression of an organ from the query dataset and that organ in the reference dataset, compared to the correlation of the organ with second-highest correlation in the reference:

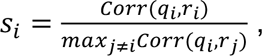

where *q*_i_ and *r*_i_ are the average expression of organ *i* in the query, co-embedded scRNA-seq or seqFISH datasets and reference, deeply profiled endoderm scRNA-seq atlas. The average significance of the scRNA-seq dataset is <u>*s*</u> = 2.76, while for the seqFISH gut tube it is only <u>*s*</u> = 1.76.

We pooled the empirically measured (seqFISH) and imputed (scRNA-seq) COVET matrices together, visualized them with UMAP and clustered them with PhenoGraph^29^ using the AOT metric (**Fig. 4a**). To assign organ labels to COVET clusters, we quantify the overlap between COVET cluster and scRNA-seq organ classification. Specifically, we calculate the ratio of cells from every organ assigned to each COVET cluster and find a strong concordance between the two cell labelings, with most organs enriched in only a single, distinct COVET cluster (**Supplementary Fig. 6b**), as apparent from the COVET UMAP (**Fig. 4b**). Using the COVET co-embedding and clustering, we could classify the seqFISH endoderm cells into distinct organs and map these back to the seqFISH image (**Fig. 4c**), where organs indeed mapped to their expected ordering and location:

- Thymus/thyroid cells fell into the most anterior COVET cluster CC0, with 75% of the compartment being assigned to it. Remaining cells were assigned almost exclusively to the spatially proximal CC1 (17%).
- 52% of dorsal lung cells were assigned to COVET cluster CC1; most remaining cells (36%) fell into cluster CC0.
- Ventral lung cells were assigned to COVET cluster CC2 (62%); remaining cells were assigned to CC1 (12%) or CC3 (19%), both highly related to CC2.
- Liver cells were almost all (94%) assigned to COVET cluster CC2.
- Pancreas cells were enriched in COVET cluster CC3 (58%); the majority of remaining cells (26%) were placed in the related cluster CC2.
- Small intestine cells were primarily in posterior COVET clusters CC4 (42%) or CC5 (49%).
- Colon cells were entirely in one of the posterior COVET clusters (CC4-CC7), with most in either CC6 (36%) and CC7 (43%).

We can also make direct use of gene-gene patterns in the ENVI predicted COVET matrices of the scRNA-seq data to elucidate patterns of niche specific gene expression, beyond endoderm cells. We demonstrate this by comparing between the dorsal and ventral lung regions using distinctly covarying groups of genes in the average COVET matrices of each comportment. We compute the AOT average for a set of COVET matrices by calculating the matrix-square of mean of their MSQR:

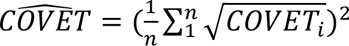

We perform hierarchical clustering on the average COVET matrices of the dorsal and ventral lung scRNA-seq and find groups of genes that uniquely co-cluster in each matrix. The genes *Dlk1*, *Gata4*, *Gata5*, *Aldh1a2* and *Foxf1* all covary in the ventral lung COVET but not in the dorsal one, while the genes *Tagln*, *Six3*, *Thbs1*, *Nkx2-3*, and *Tbx1* have the opposite pattern (**Fig. 4d**). We visualize the expression of the COVET gene sets by plotting their average expression in cells near the anterior gut tube and find that the ventral niche genes are enriched in the splanchnic mesoderm while dorsal niche genes localize to brain and cranial mesoderm (**Fig. 4e**). Gut tube cells themselves are not colored to avoid confounding the visualization with their expression. As the splanchnic mesoderm is ventral to the gut and the brain and cranium are dorsal, the differentially covarying genes in the COVET matrices allow us to reconstruct each lung compartment’s spatial context.

We tried to perform similar analysis with other integration methods which, unlike ENVI, do not model cellular environments, namely gimVI, Tangram and Scanorama^49^. With each method, we attempted to label the seqFISH gut tube cells using the scRNA-seq organs as reference (**Supplementary Fig. 6c**). Only ENVI explicitly predicts spatial context of the scRNA-seq data, so for a comparison between methods, we use their combined latent or learned mapping to label the seqFISH gut tube cells into organs and qualitatively evaluate how each compared to known pattern of organogenesis. For methods with combined embeddings (ENVI COVET, gimVI and Scanorama), we label using a kNN classifier, which assigns each seqFISH gut tube cell an organ based on the most common label of its *k* = 5 scRNA-seq gut tube neighbors in the combined space:

- ENVI: To label seqFISH cells, we fit an AOT metric kNN classifier (*k* = 5) on the scRNA-seq ENVI COVET matrices and their organ labels, and applied the classifier to the seqFISH COVET matrices. Note that this is a different approach than the clustering-based approach used above to predict spatial context for the scRNA-seq gut cells. Here, we are trying to assign labels from the scRNA-seq to the spatial data, so we employ a discrete classifier.
- gimVI: We trained gimVI on the full embryogenesis scRNA-seq and seqFISH datasets using the same default parameters as in the *Benchmarking imputation* section above (10 latent dimension, 200 epochs, NB for spatial and ZINB for single-cell). We subset the learned latent embedding of scRNA-seq and seqFISH modalities to only the gut tube cells from each dataset, and similarly learned a kNN classifier (*k* = 5) from the single-cell latent and organ assignment, using it to predict labels on the spatial latent.
- Scanorma: Scanorama is a general batch-correction method, not specifically designed to integrate spatial and scRNA-seq data. It uses mutual nearest neighbors to directly correct the gene expression count matrix and remove batch effect. Following the instructions in *scanpy* (scanpy.readthedocs.io), we applied Scanorama to the scRNA-seq and seqFISH gut tube cells, where the scRNA-seq only includes the seqFISH genes. Specifically, we concatenated the seqFISH and scRNA-seq gut tube cells in a single dataset, calculated the top 50 PCs based on logged expression, and trained Scanorama, with the *batch* parameter set to integrate across data modality. With the corrected count matrices, we trained a kNN (*k* = 5) classifier on scRNA-seq labels and assigned organs to the seqFISH gut tube cells.
- Tangram: Using the parameters laid out in *Benchmarking imputation* (1,000 epochs, uniform density prior, ‘cells’ mode), we trained Tangram to learn a mapping matrix from scRNA-seq to spatial data. We subset the Tangram matrix to the mapping from scRNA-seq gut tube to seqFISH gut tube cells and re-normalized the columns to sum to 1. We transferred organ labels using Tangram’s *project_cell_annotations* function, which uses the subsetted mapping matrix to calculate the probability of each organ being assigned to each seqFISH gut tube cell, and we labeled according to the most probable organ.

### AP polarity of developing spine and NMP cells

On spinal cord and neural mesoderm progenitor (NMP) cells, which span the embryo AP axis, we found that the first COVET DC and primary FDL axis were highly congruent with AP polarity (**Fig. 5a**), as apparent in seqFISH data (**Fig. 5b**). DCs were computed via eigendecomposition of the Laplacian of the kNN (*k* = 30) graph in COVET space, where the graph is created from COVET matrices from both seqFISH (measured) and scRNA-seq (ENVI-predicted) spine and NMP cells. To quantify the agreement between pseudo-AP and true AP on the seqFISH data, we computed DCs of the seqFISH spine and NMP cells based on their spatial coordinates, and found that the first DC visually aligns with AP polarity (**Supplementary Fig. 7a**). The seqFISH coordinate DC (true-AP) and COVET DC (pseudo-AP) were highly congruent, having a Pearson correlation of *r* = 0.86.

Using gimVI and Scanorama, we attempt to similarly reconstruct a pseudo-AP axis from DC analysis of their latent spaces, and study how well it matches the true AP axis:

- gimVI: We used the gimVI model trained on the complete embryogenesis datasets, and subset the learned gimVI combined latent to only the spine and NMP cells from the spatial and scRNA-seq datasets. We calculate the top 3 DCs from both latent embeddings and find that the 2nd DC had the highest correlation with the true AP polarity on the seqFISH spine and NMP cells *r* = 0.76
- Scanorama: We applied Scanorma as described in *Spatial organization of emerging organs* to produce integrated count matrices of the seqFISH and scRNA-seq spine and NMP cells. We computed DCs from the combined Scanorama-corrected scRNA-seq and spatial datasets and found that the third DC had highest correlation with seqFISH true AP *r* = 0.70.

Although we selected the DC that best matches the true AP for gimVI and Scanorama, both methods produced spine and NMP cells in the posterior with low DC values, unlike the consistent trend in ENVI (**Supplementary Fig. 7b**). We note that since DC order is arbitrary, we reversed any DC which had negative correlation with the true AP. Tangram was excluded from this analysis as it does not calculate a combined embedding from which we can recover a pseudo-AP axis.

To compare how accurately the pseudo-AP from ENVI, gimVI and Scanorama map on the scRNA-seq spinal cells, we ordered the cells based on their pseudo-AP values and visualized the expression of canonical AP markers *Rfx4*^55^ (anterior), *Hoxaas3*^54^ (posterior) and *Hoxb7*^56^ (posterior) along the predicted axes (**Fig. 5e**). Gene expressions were logged and z-scored with ordered profiles being smoothed with a first-order Savitzky-Golay filter with window size 128 for visual clarity.

We next quantify the quality of the pseudo-AP each method predicted for the scRNA-seq spinal cells. For each pseudo-AP axis of the scRNA-seq data, we calculate its correlation with the (logged) expression of 5 known posterior genes (*Hoxaas3, Hoxb5os*^54^*, Hoxb9*^53^*, Hoxb7* and *Tlx2*^78^) and anterior genes (*Foxa3*^79, 80^*, Hoxd3*^52^*, Hoxa2*^81–83^*, Rfx4* and *Hoxd4*). The quantification (**Supplementary Fig. 7c**) recapitulates the qualitative results implied by the pseudo-AP ordered expression profile (**Fig. 5e**). We see that ENVI pseudo-AP is consistently strongly correlated with posterior markers and anti-correlated with anterior markers, while gimVI consistently reverses the pattern, and Scanorma, albeit correctly predicting polarity trend, scores lower than ENVI.

### Mapping cortical depth of dissociated neurons with MERFISH

We used the Brain Initiative Cell Census Network’s 254-gene MERFISH primary motor cortex dataset^38^, along with its matching scRNA-seq reference^57^ to demonstrate ENVI in a tissue-wide context. To prevent dataset specific batch effects from confounding our integration results, we selected only the first sample from the scRNA-seq atlas (labeled ‘L8TX_181211_01_B01’). We kept only the cells with cell type annotations and removed any labeled as doublets or low quality, leaving 7,416 cells in the dataset. We also removed any genes that both i) appear in less than 5% of cells and ii) are not in the MERFISH panel. We similarly used only the first sample from the BICCN MERFISH data (‘dataset1_sample_1’) which included 18,516 cells from 6 individual slices of the motor cortex. We removed cells without a cell type label and the genes *Crispld2* and *Igf2* from the MERFISH data as they were not included in the scRNA-seq and avoided any additional pre-processing steps. Both spatial and scRNA-seq datasets were labeled into neuronal and non-neural cell types. For brevity and consistency between datasets, we relabeled the MERFISH GABAErgic neurons from ‘Sst-chodl’ to ‘Sst’ and collapsed the ‘PVM’, ‘macrophage’ and ‘microglia’ labels to ‘microglia’.

We benchmarked ENVI, Tangram, gimVI and NovoSpaRc imputation on MERFISH motor cortex, using fivefold cross validation (see *Benchmarking imputation*). Imputation quality was assessed with MSSI and Pearson correlation, and ENVI outperformed all methods with strong statistical significance, according to one-sided relative t-test. Additionally, we visualized the ground-truth and predicted expression of two genes with clear localization in distinct cortical layers: *Prdm8* and *Satb2*. On one of the MERFISH slices, we plotted the (logged) ground-truth and imputed (ENVI, Tangram and gimVI) expression. For both genes, we found that only ENVI visually matches true expression, while Tangram only predicted their expression to a subset of the cells that do, and gimVI overestimated their abundance (**Fig. 3g**).

To map scRNA-seq glutamatergic neurons to their cortical depth, we co-embedded the ENVI-imputed COVET matrices from the scRNA-seq data and the measured COVET matrices from the MERFISH data constructing a KNN graph (*k* = 100) using the AOT distance. We calculated COVET DC and FDL. We find that the first COVET DC (pseudodepth) matched the cortical depth of the MERFISH compartment, as visualized on the MERFISH slice (**Fig. 6b**). We validated the inferred cortical depth using this approach with the cortical depth labeling provided in Yao et al.^57^, deduced from known molecular markers to cortical depth. Indeed, pseudodepth derived from ENVI predicted COVET matrices matches the provided scRNA-seq cortical depth annotation (**Fig. 6c**).

We similarly trained gimVI and Scanorama to integrate between the two datasets to find correlates with the cortical depth.

- gimVI: We trained the default model with the default parameters described *Benchmarking Imputation* on the spatial and single-cell datasets. We subset the learned latent to only include the glutamatergic neuron compartment of the two datasets and calculated the top 3 DCs.
- Scanorma: Scanorama is trained only on the glutamatergic neurons of the MERFISH and scRNA-seq datasets, with the same default configuration as in the *AP polarity of developing spine cells* section. We computed the top 3 DCs from the integrated expression profiles of the two datasets.

We visualized the DC from each method (ENVI, gimVI and Scanorama) on one MERFISH slice (**Supplementary Fig. 8a**), and found that qualitatively, only the first ENVI-derived DC accurately recapitulates cortical depth. To explicitly quantify the pseudodepth derived from each method, we first assigned each glutamatergic neuron in the MERFISH data a true-depth value using DC analysis of their spatial coordinates. As the MERFISH data is divided into slices, we calculated separate coordinate DCs from each. Visually, the second DC best matched cortical depth across all 6 slices, and the DC was reversed if it was opposite to the true depth, as DC direction is arbitrary. We combined the true depth of each individual slice to one scale for the entire dataset by normalizing (min-max) each slice separately to values between 0 and 1.

Pseudodepth of gimVI and Scanorama is defined as the DC most highly correlated with the true depth, which was represented by the first DCs for both algorithms. We calculated the Pearson correlation between MERFISH pseudodepth derived from ENVI, gimVI and Scanorma; across all slices (**Supplementary Fig. 8b**). We also compared how each pseudodepth axis aligned with the layer-related labels of the glutamatergic cells for both the scRNA-seq and MERFISH datasets. We calculated the average pseudodepth of each method across cells in each cortical layer (L2/3, L5, L5/6 and L6) for both. Only ENVI pseudodepth correctly scales from L2/3 to L6, with both gimVI and Scanorama incorrectly swapping the order of layers (**Supplementary Fig. 8c**).

We ordered the scRNA-seq glutamatergic neuron cells by their ENVI-inferred pseudodepth (**Fig. 6d**), only including genes expressed in more than 5 glutamatergic neurons and that are within the union of the top 8,192 HVGs and MERFISH genes. Genes were ordered according to the pseudodepth at which they peak. As the expression of a specific gene can be noisy, to determine the pseudodepth peak of a gene we ordered the cells and smoothed their expression along the axis with a 32-length, running average filter. We assigned pseudodepth to a gene based on the pseudodepth of the cell (smoothed) where it is most highly expressed, and found cortical gradients based on the pseudodepth-ordered gene expression heatmap (logged and z-scored). For clarity, we slightly smoothed the resulting heatmap with a 4-length running average filter.

To capture the transition between cortical layers in the scRNA-seq glutamatergic neurons, we looked to gene sets which describe the programs activated in each distinct layer. The Allan Brain Atlas have identified differentially expressed genes (DEG) associated with each cortex layer (L2/3, L4, L5 and L6). We use these to determine a gene set for each layer, by i) taking the top 400 ISH upregulated (log-fold change) probes from the Allan Brain Atlas, ii) collapsing probes that target the same gene, and iii) filtering genes that are not highly variable (**Fig. 6d**). To derive a final gene set, for genes present in more than one set, we retained only the expression in the layer where their log-fold change is highest to get only distinct genes that represent each layer. Finally, we only kept the top 64 genes (based on log fold change) in each set (**Supplementary Table 3**). We marked the pseudodepth of the genes (**Fig. 6d**, right), and found that they are appropriately ordered from L2/3 to L6, including a distinct L4 compartment, further supporting the suggestion^59^ that there are dedicated L4 neurons in the motor cortex.

We plot the average expression (z-scored, logged) of each gene set in the scRNA-seq glutamatergic neuron, ordered by their pseudodepth (**Fig. 6e**). Ordered average expression profiles are smoothed by a 128-length running average. Here too we see that there is a distinct region of cells enriched of L4 markers, immediately after the L2/3 and before L5 regions. We similarly order MERFISH glutamatergic neurons by their true- depth, and plot the (z-scored, logged) average expression of each cortical layer gene set, subsetted to only include the genes in the MERFISH assay, smoothed with a 128-length running average. We see that there is no clear separation between L4 and L2/3 related expressions (**Fig. 6f**).

---

**Supplementary Figure 1.**
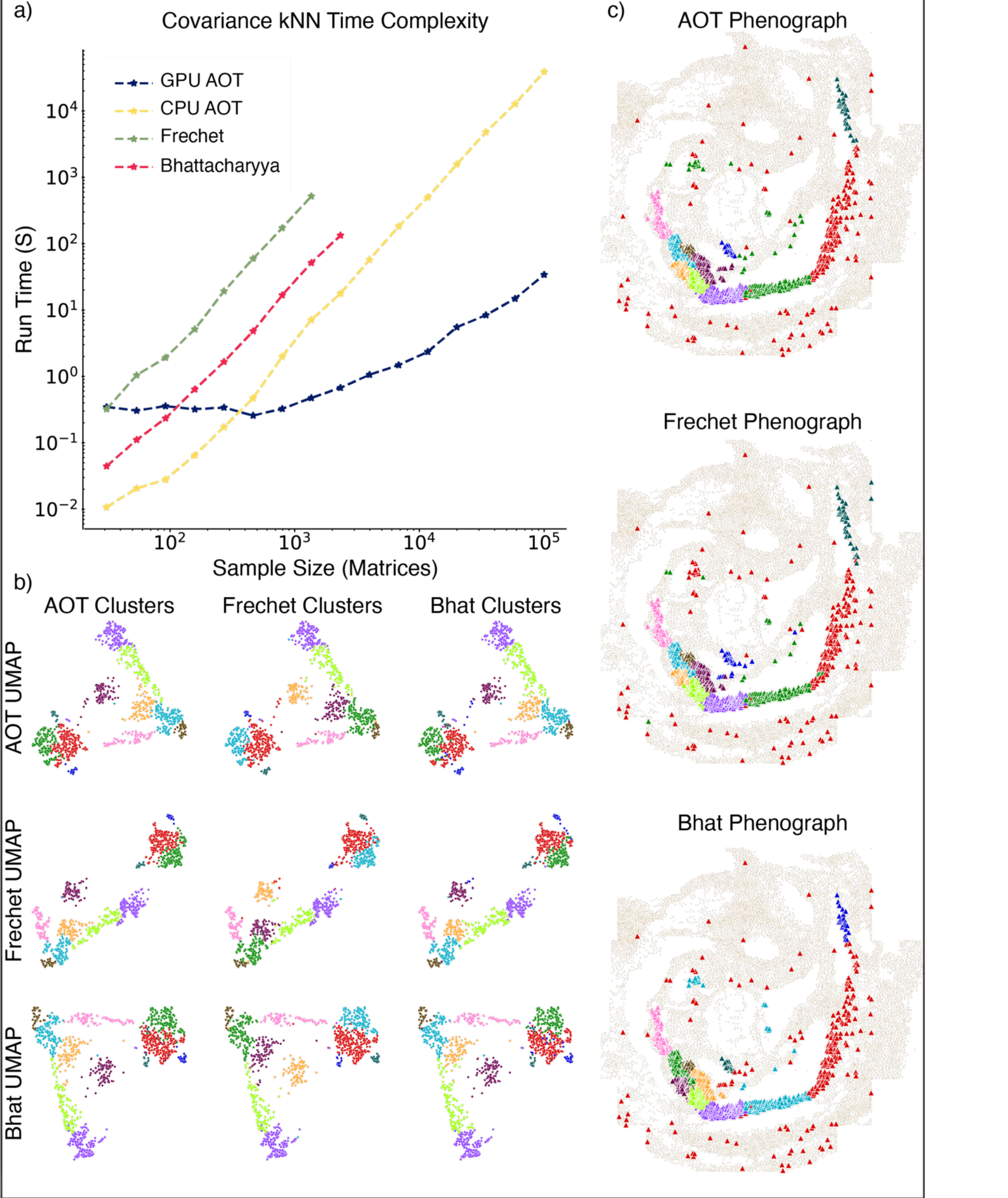
Approximate optimal transport (AOT) yields similar results to optimal transport and Bhattacharyya distance, but more efficiently. **a**, Run times for computing the kNN graph between sets of randomly generated covariance matrices at various sample sizes. Fréchet and Bhattacharyya run times are not shown for samples larger than 4,000 cells due to out-of-memory error on a 768-GB, 64-core computing cluster. **b**, COVET UMAP embeddings and PhenoGraph clustering of seqFISH splanchnic mesoderm by different metrics, colored by PhenoGraph clusters of each. **c**, seqFISH data from splanchnic mesoderm, colored by PhenoGraph clustering of COVET matrices according to each distance metric. Bhat, Bhattacharyya.

**Supplementary Figure 2.**
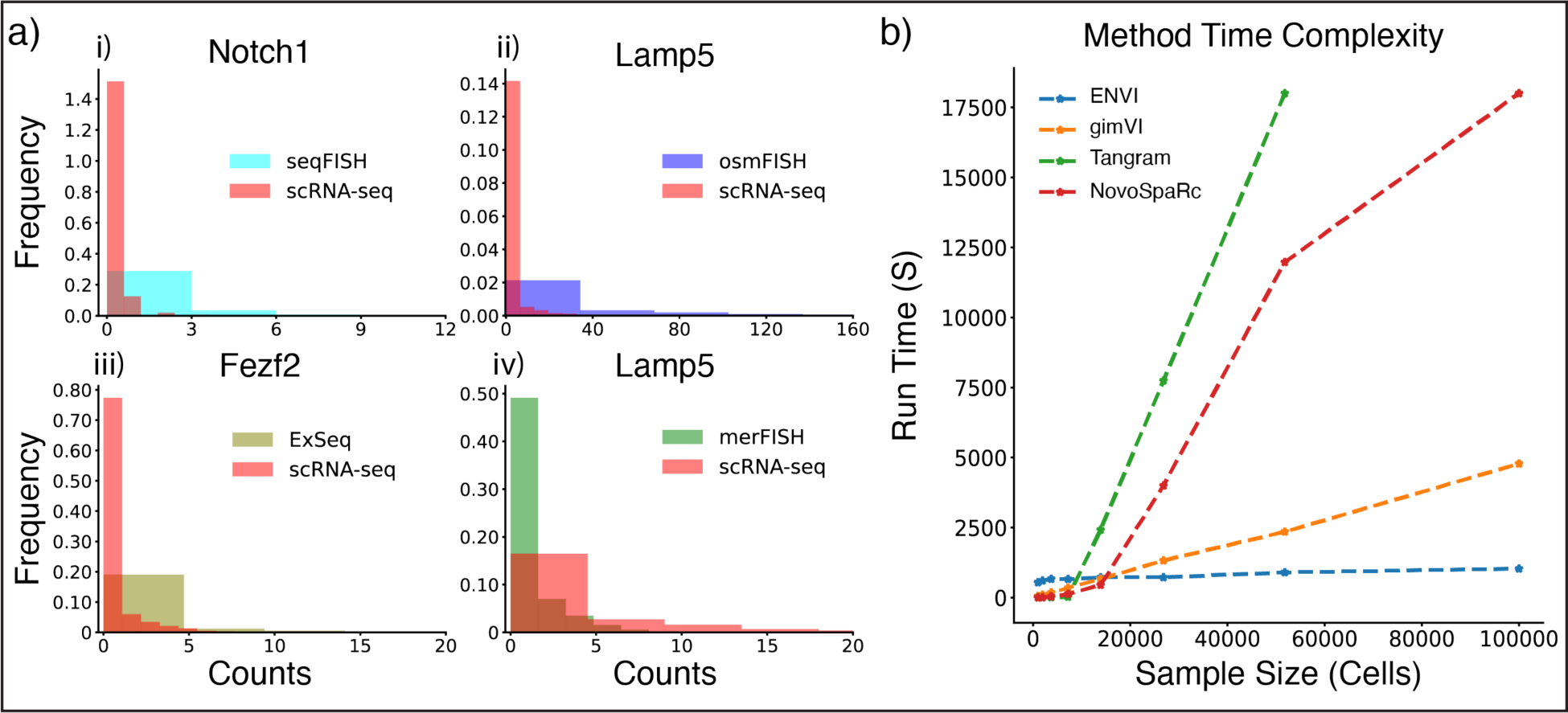
Differences between spatial and scRNA-seq data and run time for different integration methods. **a**, The expression of three genes in four spatial datasets exhibits very different distributions from their complementary scRNA-seq data. **b**, Run time of ENVI, gimVI, Tangram and NovoSpaRc on integrating simulated scRNA-seq and spatial datasets of growing sizes. Run time for each method was capped at 5 hours (18,000 seconds) before the program was manually stopped.

**Supplementary Figure 3.**
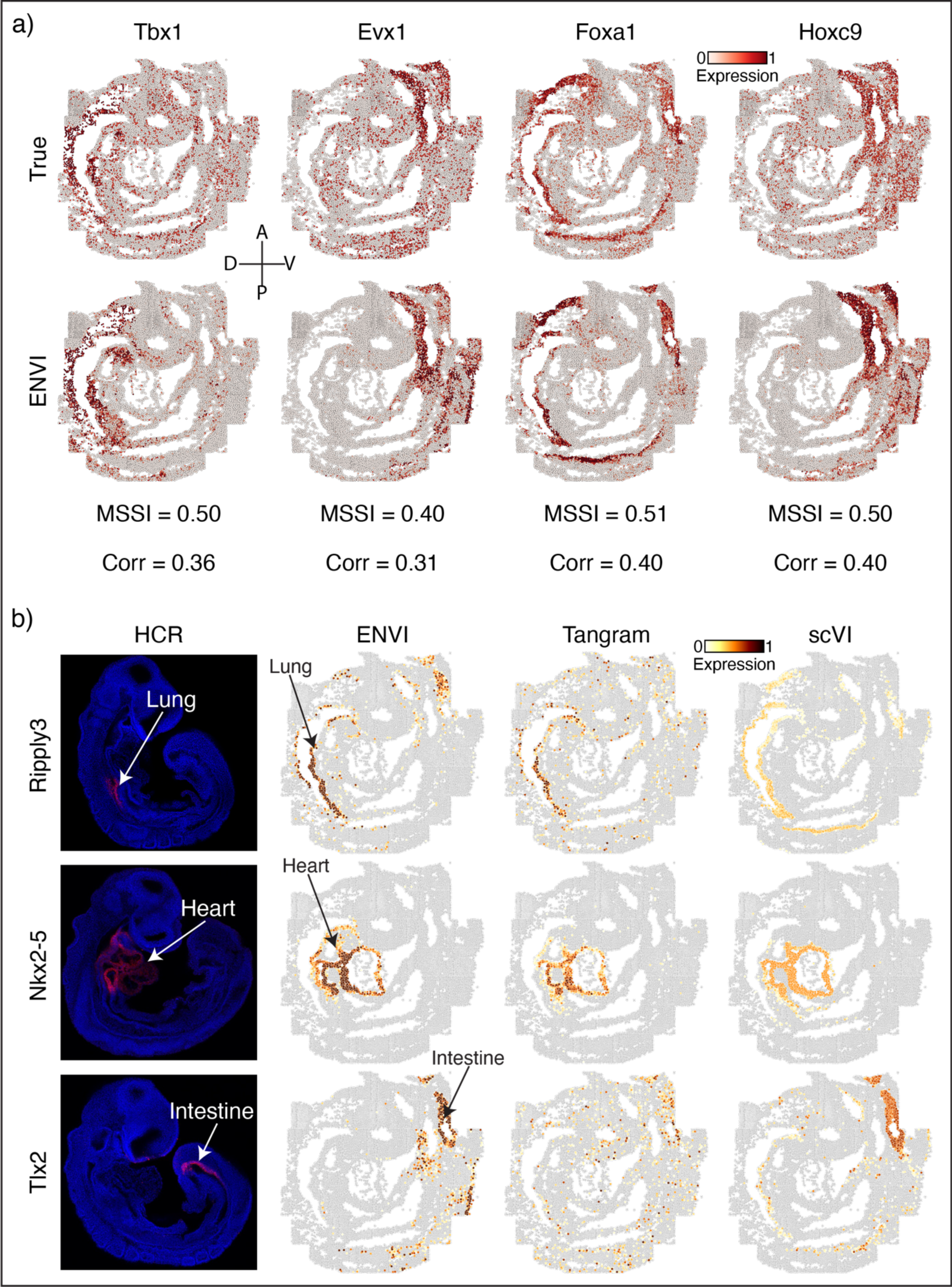
ENVI more accurately infers embryogenesis genes missing from a seqFISH panel compared to other methods. **a**, Imputed expression of withheld genes from the embryogenesis dataset (bottom) compared to true (measured) expression (top), with corresponding MSSI and Pearson correlation reconstruction scores below. **b**, HCR images of *Ripply3*, *Nkx2-5* and *Tlx2* and their imputation values according to ENVI, Tangram and gimVI. Organs marked by each gene are noted on the HCR and seqFISH images.

**Supplementary Figure 4.**
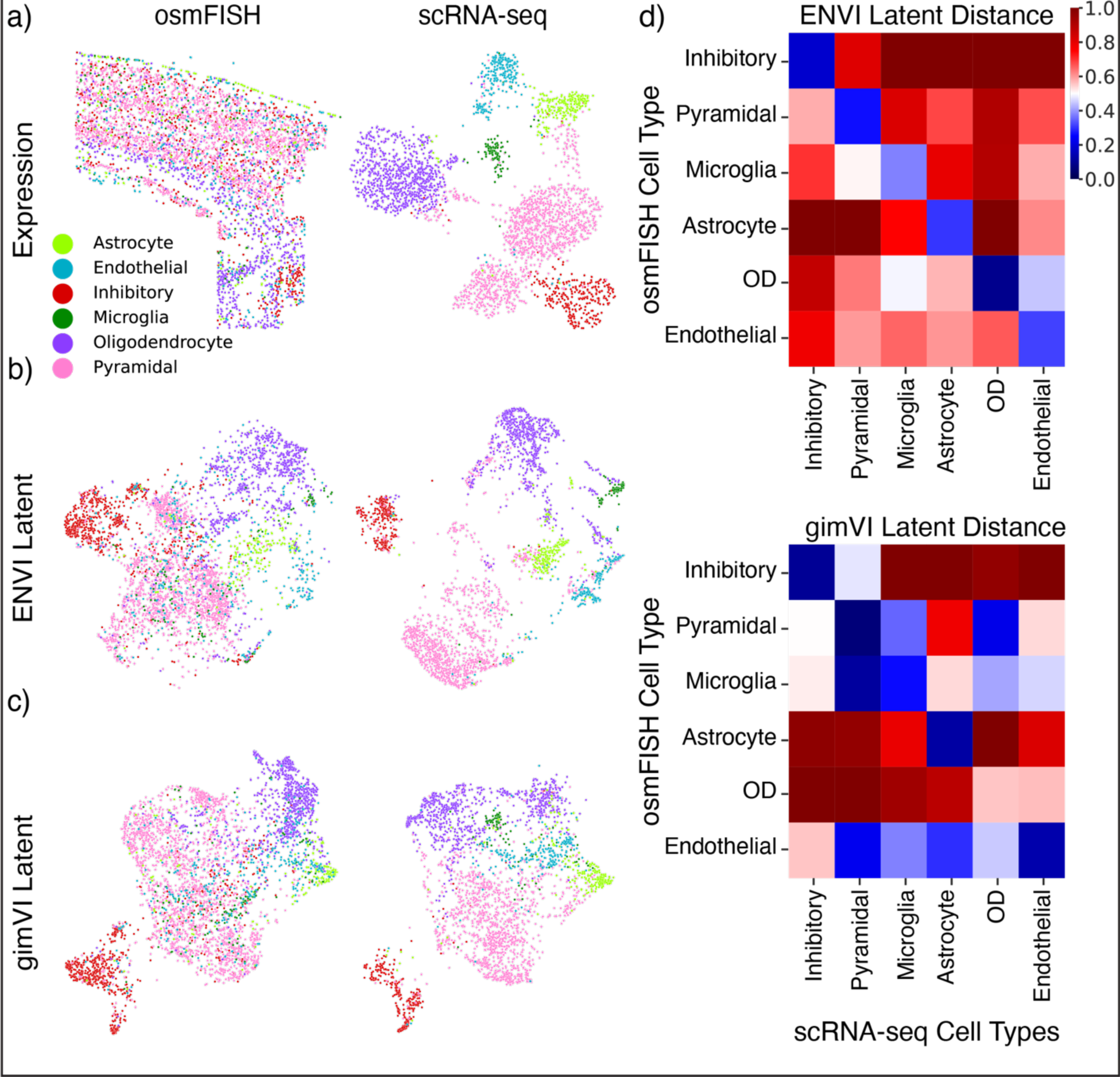
ENVI can integrate between a 33 gene osmFISH panel and scRNA-seq assay of the somatosensory cortex. **a**, osmFISH with segmented cells and UMAP visualization of scRNA-seq datasets of the mouse somatosensory cortex, colored by cell types as annotated in Codeluppi et al^3^. **b**, UMAP visualizations of the ENVI integrated latent embedding of the osmFISH and scRNA-seq modalities, colored by cell types as in **a**. **c**, Same as **b**, but with latent embeddings from gimVI. **d**, Normalized distance between the center-of-mass of each cell type according to the ENVI and gimVI latent embeddings.

**Supplementary Figure 5.**
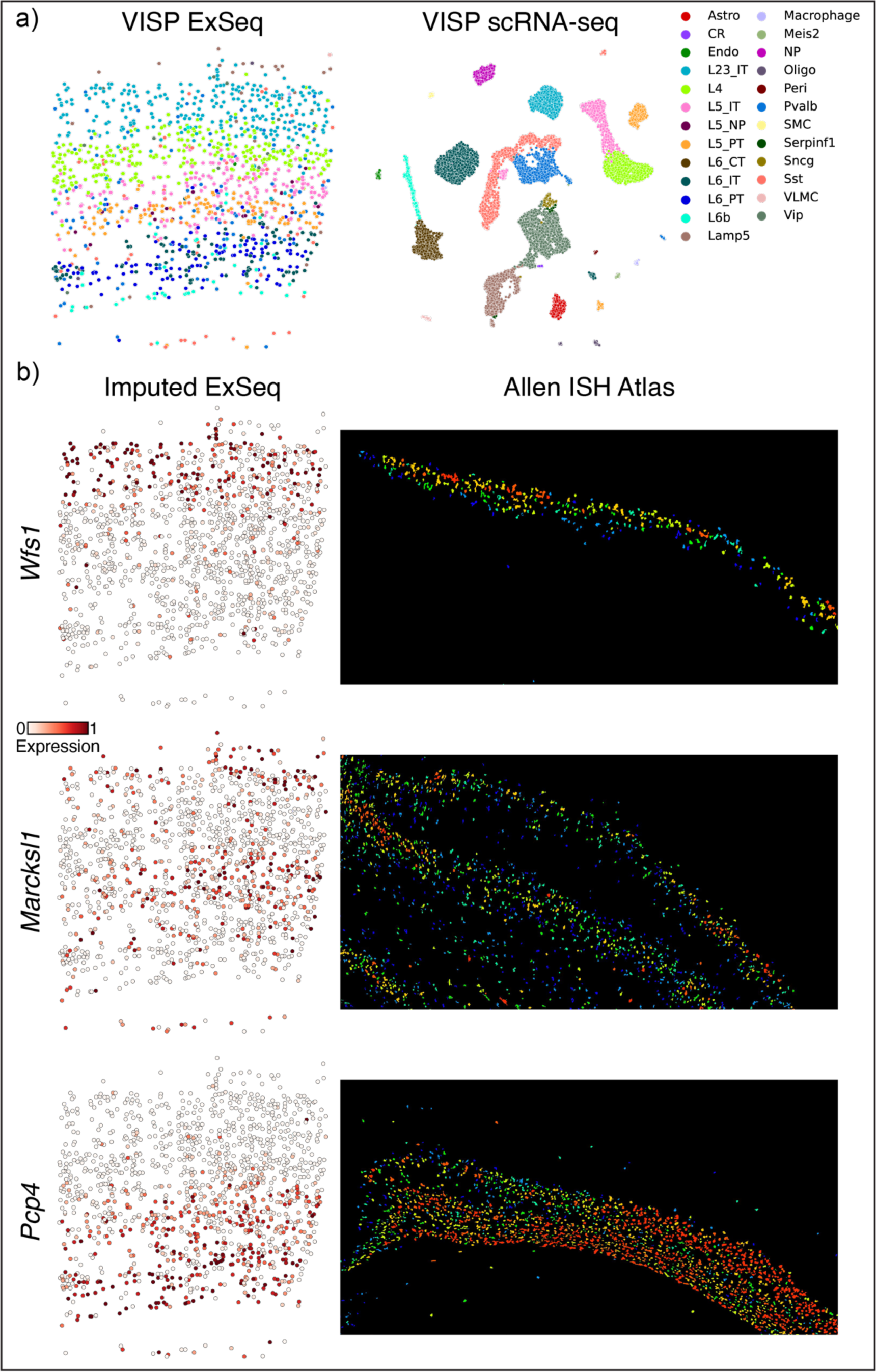
ENVI generalizes to imaging technologies beyond in situ hybridization (ISH). **a**, Spatial ExSeq (VISP^37^) data of the visual cortex (left) and the complementary scRNA-seq data, visualized by UMAP (right), colored into cell types. **b**, ENVI- imputed expression of the unimaged cortical markers *Wfs1*, *Marcksl1*, and *Pcp4* (left) and ISH imaging of the visual cortex from the Allen Brain Atlas (mouse.brain-map.org) (right).

**Supplementary Figure 6.**
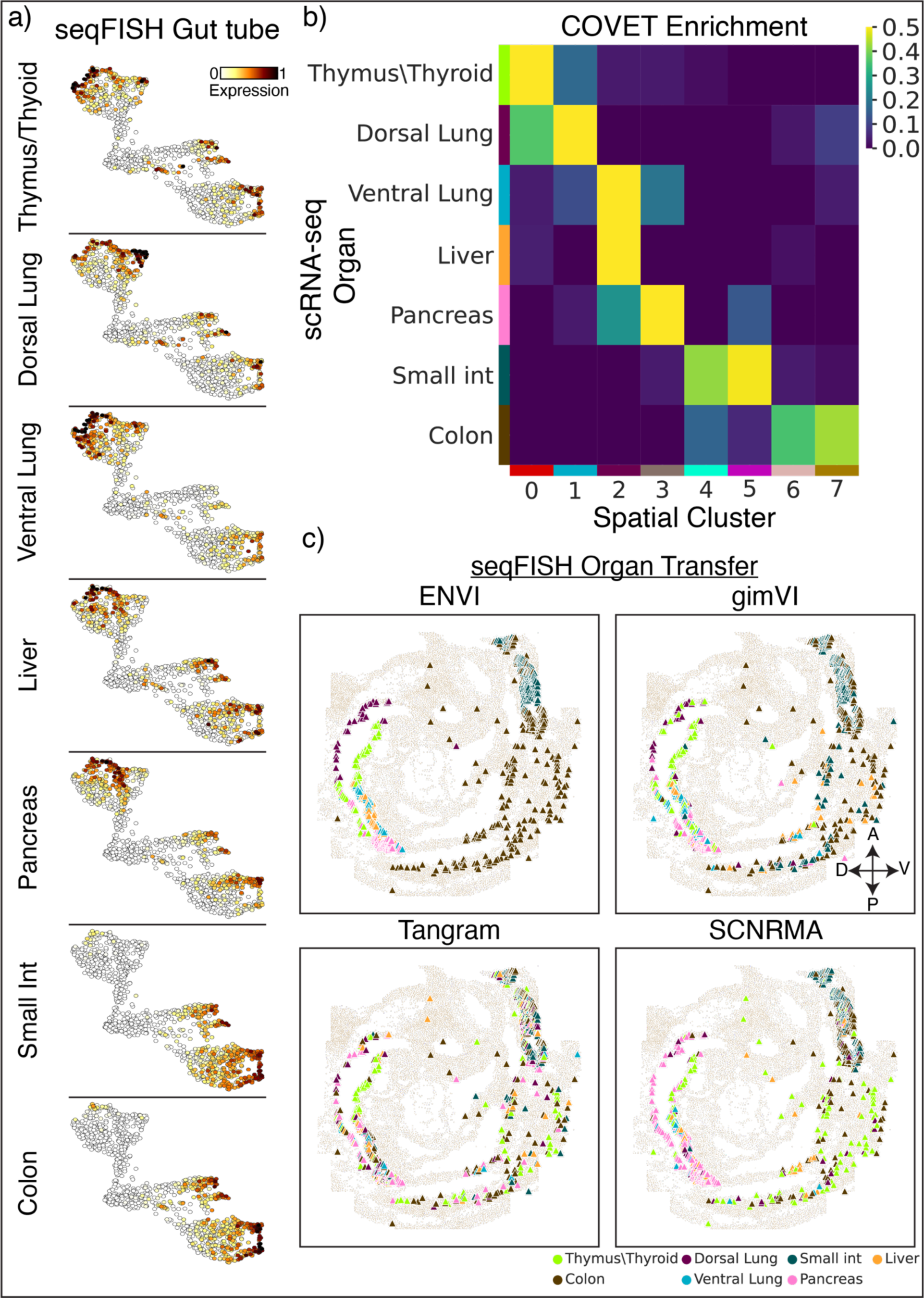
Spatial orientation of gastrulation from ENVI. **a**, UMAP of gut tube cells from seqFISH expression data^35^, colored by average expression of reference gene sets. **b**, Proportion of scRNA-seq gut tube cells in each organ (row) which fall in each COVET cluster (columns). **c**. Assignment of developing organs to seqFISH gut tube cells via ENVI COVET space, gimVI latent space, Tangram cell-type mapping and Scanorama ’integrated’ expression profiles, using the scRNA-seq labeled gut as reference.

**Supplementary Figure 7.**
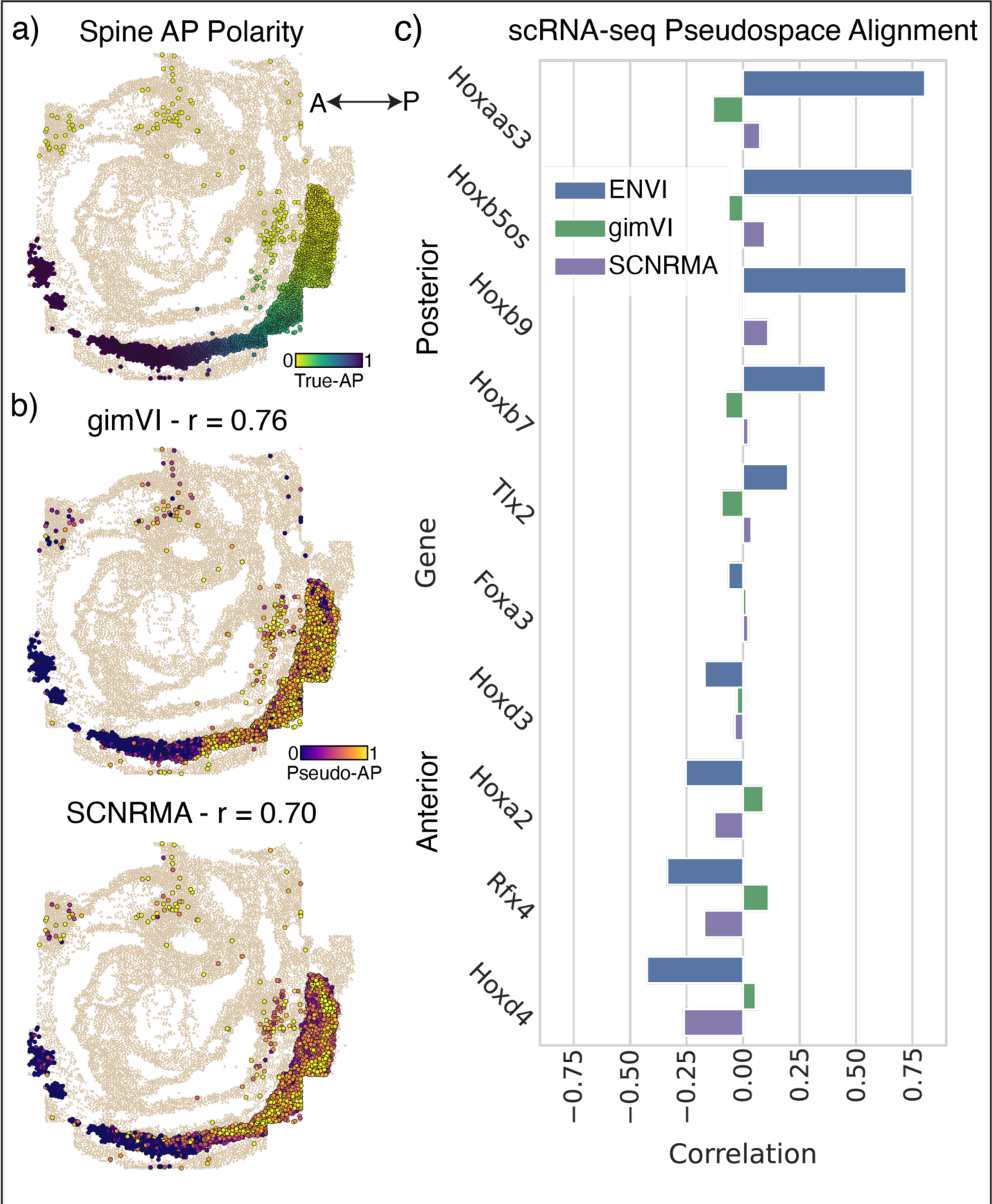
ENVI can reliably recover the AP axis during spine development. **a**, Spine and NMP cells from seqFISH data, colored by AP polarity calculated from the first DC of their spatial coordinates. **b**, Pseudo-AP of seqFISH spine and NMP cells from DC analysis of gimVI and Scanorama. Values denote the Pearson correlation with the true AP axis. **c**, Pearson correlation of ENVI COVET, gimVI and Scanorama pseudo-AP of spine and NMP scRNA-seq cells, for five canonical posterior markers (higher is better) and anterior markers (lower is better). Pseudo-AP axis is based on the diffusion component best aligned with true depth (DC 1, 2 and 3 for ENVI, gimVI and Scanorama, respectively).

**Supplementary Figure 8.**
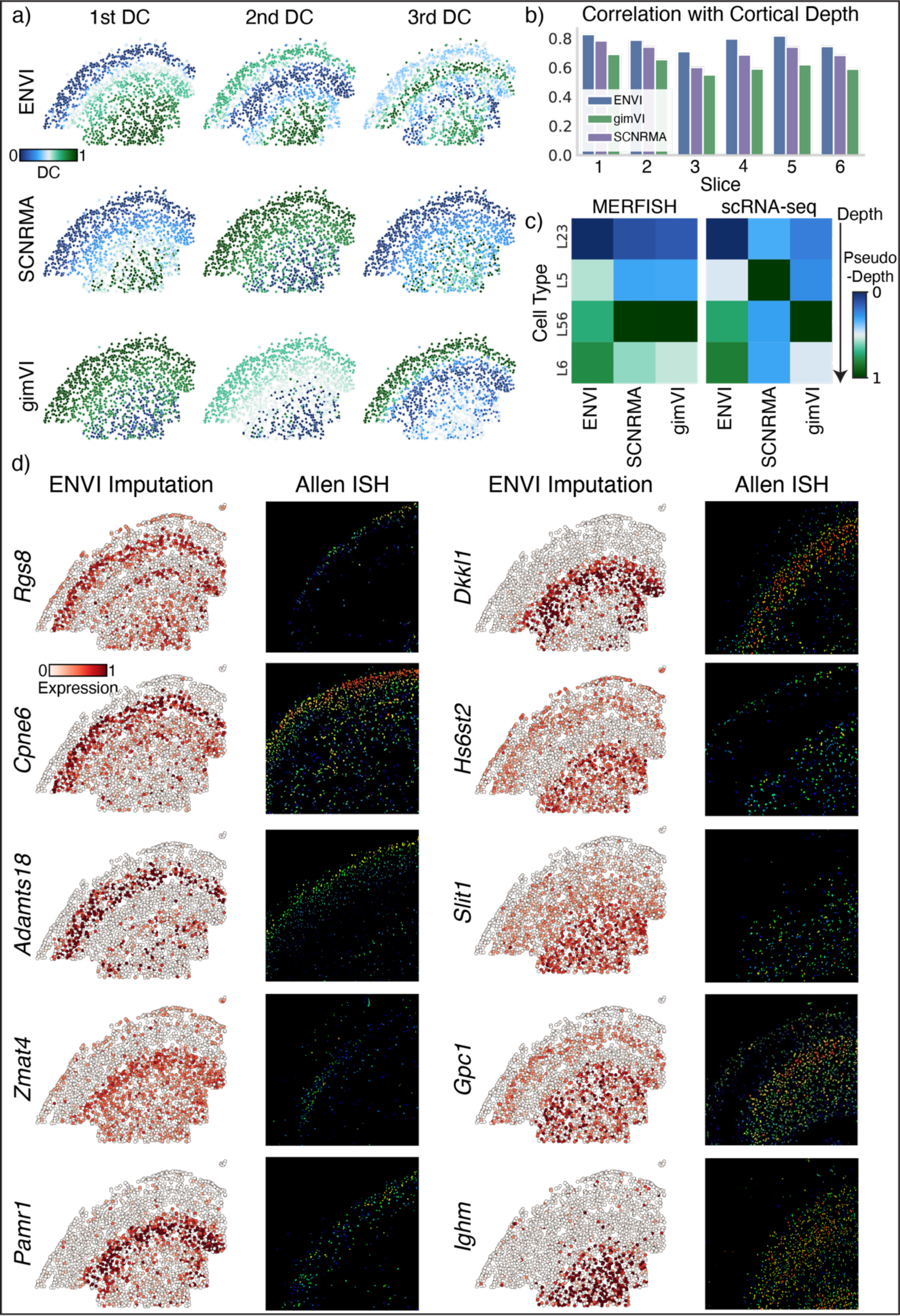
ENVI extends MERFISH panels to thousands of genes and predicts cortical depth of disassociated data. **a**, Glutamatergic neurons from the MERFISH motor cortex dataset^38^, colored by the top three DCs from ENVI COVET, gimVI and Scanorama (SCNRMA). DC analysis for all methods was performed on the MERFISH and scRNA-seq co-embedding. **b**, Quantitative comparison of methods. Pearson correlation of pseudodepth with true cortical depth for glutamatergic neurons for each tissue slice in the MERFISH dataset. True depth for each slice was defined as the second DC of the spatial coordinates of the cells. Pseudo-Depth of each method was chosen as the DC most aligned with true depth (1^st^ DC for all methods). **c**, Average z-scored pseudodepth according to each integration method of the glutamatergic neuronal subtypes, grouped by cortical layer. **d**, ENVI imputation of genes highlighted in Fig. 6d, and projected onto the MERFISH data, with corresponding ISH expression within the motor cortex from the Allen Brain Atlas (mouse.brain-map.org).

